# Effects of introgressed Neanderthal alleles on present-day brain morphology

**DOI:** 10.64898/2026.04.14.718380

**Authors:** Roberta Zeloni, Alessandro Amaolo, Adeline Morez Jacobs, Ettore Zapparoli, Yara Akl, Mahan Shafie, Emilia Huerta-Sanchez, Fabrizio Pizzagalli, Paolo Provero, Luca Pagani, Davide Marnetto

**Affiliations:** Dept. of Neurosciences “Rita Levi Montalcini”, University of Turin, Torino, Italy; Dept. of Biology, University of Padova, Padova, Italy; Epigenetics and Neurobiology Unit, European Molecular Biology Laboratory (EMBL) Rome, Monterotondo, Italy; Department of Genetics, Trinity College Dublin, Ireland

## Abstract

Neanderthal introgression contributed a small fraction of genetic variants to present-day non-African genomes. While differences in cranial globularity between Neanderthal and modern humans are well documented from endocasts, the phenotypic consequences of these introgressed alleles can illuminate otherwise inaccessible genetically divergent brain structures.

We analyzed 370 MRI-derived brain traits—including cortical and subcortical regional measurements, cortical folding metrics, diffusion tracts—in nearly 40,000 UK Biobank participants. To quantify the impact of Neanderthal ancestry, we intersected trait-associated loci with Neanderthal-derived variants identified from introgressed segments imputed in the same subjects.

Low-frequency introgressed variants were depleted for detectable effects on brain phenotypes, whereas common introgressed variants showed no comparable depletion. Conversely, Neanderthal deserts were consistently enriched for functional effects. Eight associations were fine-mapped to Neanderthal-derived variants: one locus near the gene DAAM1 was especially prominent across multiple traits, including opposite effects in the cuneus and precuneus mediated by introgressed regulatory variants. Genome-wide directional alignment of Neanderthal effects was limited but became evident when focusing on suggestive loci: frontal and parietal areas were the most consistently affected traits, though not in a direction that obviously mirrors known modern-archaic morphology divergence. Several of these loci also influenced neuropsychiatric traits, with detectable polygenic consequences against schizophrenia and towards major depression, linking neuroanatomical and neuropsychiatric impact of Neanderthal introgression.

These findings suggest that while introgressed alleles affecting divergent neuroanatomy between modern humans and Neanderthals were largely purged, a subset of tolerated alleles continues to shape human brain morphology and mental health.

## Introduction

Around 40 thousands of years ago (kya), ancestors of anatomically modern humans (AMH) experienced a period of interbreeding with Neanderthals, an archaic human group that diverged from the modern human lineage around 600 kya^1,2^. As a consequence, present-day non-African humans harbor on average 2% introgressed haplotypes, scattered around to cover about 35% of the human reference genome^3–6^. Neanderthal introgressed segments have become part of the genetic pool of non-African modern humans and have contributed to their differentiation. Although some examples of adaptive introgression from archaic humans have been identified^7–10^, most introgressed haplotypes are expected to be neutral, or even fitness-reducing due to a) critical demographic conditions of late Neanderthals leading to accumulation of deleterious variants^6,11^ and b) incompatibility with the rest of the modern human genome^6,12^. Indeed, selection against functional introgressed variants has been demonstrated^6,12–14^, and some regions, termed archaic deserts, which make up to 7-40%^5,6,15,16^ of the modern human genome, are devoid of recognizable Neanderthal introgression.

Previous research has shed light on the influence of remaining Neanderthal-derived variation on present-day phenotypic diversity^17–23^, identifying phenotypic domains significantly shaped by introgressed alleles and the variants driving those associations. Among others, behavioural and neuropsychiatric traits emerged as especially impacted: depression^21^, schizophrenia and anorexia^17^ were found significantly affected; sleeping patterns, pain, mental health, smoking and alcohol consumption even showed signs of enrichment in Neanderthal-derived variants^18^. This is especially striking in a context of overall depleted impact of Neanderthal introgression to the heritability of complex traits^17,19^.

As the most iconic morphological divergence between AMH and Neanderthals is the craniofacial shape^24–26^, it is compelling to conjecture a parallelism between divergent evolution in brain morphology and a major impact on behavioural and neuropsychiatric traits.

Neanderthal’s outer brain morphology, inferred from casts of the braincase (“endocasts”), is characterized by a more elongated shape along the anterior-posterior axis, whereas AMH evolved over time a more globular shape^25,27,28^. This divergence likely develops in the perinatal period^29^ and is chiefly derived from a bulging cerebellum and parietal lobes, together with a smaller occipital lobe^25,27,28^. Parietal bulging, however, is not paralleled by a general area expansion^27,30^, suggesting rather a size increase of internal regions that displace the parietal surface. The precuneus, which is the medial component of the parietal lobe, is a potential candidate for driving parietal bulging through internal expansion due to its recent evolutionary changes^31^, but its shape cannot be directly gleaned from endocasts.

An alternative mode of exploration of Neanderthal-specific brain morphology focuses on the present-day effects of introgressed genetic variants. Applying this genetic approach Gunz et al.^27^ identified two Neanderthal-derived genetic loci associated with present-day brain globularity encompassing the *UBR4* and *PHLPP1* genes. On the other hand, Gregory et al.^32^ found positive correlations between genomic Neanderthal burden and depth of the right intraparietal sulcus and visual cortex surface (occipital lobe), cortical features that cannot be resolved from endocasts. However, the study was conducted on only 221 samples, which limited statistical power and prevented the identification of robustly associated loci. The advent of MRI databases now encompassing tens of thousands of individuals offers a substantial opportunity to redefine and expand these findings^33,34^. Singling out the brain features that diverged the most after the Neanderthal-AMH split can not only elucidate Neanderthal brain features which are not appreciable from the analysis of fossil braincases, but allow us to identify the most recent evolutionary innovations developed in our lineage. Nevertheless, a thorough large-scale analysis of the effect of archaic variants on present-day human brain morphology measured through MRI is still lacking. With this study we aim to fill this gap.

Following the imputation of Neanderthal introgressed segments in the UK Biobank, we identify single variants which diverged between Neanderthal and AMH, and have a genome-wide significant effect on brain imaging-derived phenotypes (BIDPs). To assess the selective regime acting on these variants after their introduction through introgression, we quantify the relative abundance of significant effects for Neanderthal-derived variants and Neanderthal deserts. As these traits are essentially polygenic, we investigate the genome-wide directional effect of Neanderthal introgressed variants, combining summary-based and individual-based methods. Finally, to reveal possible consequences of Neanderthal-impacted morphology features, we explore the effects of these variants on behavioural and neuropsychiatric traits.

## Results

### Probing the Neanderthal legacy in the UKBB

In order to capture Neanderthal-derived variability relevant to our work, we analyzed a set of 102 million individual Neanderthal introgressed segments inferred in 45,000 UK Biobank (UKBB) samples with available brain MRI scans and presented in Morez Jacobs et al.^35^. These segments were inferred without relying on a reference Neanderthal genome using SSTAR^36^, which identifies 50-kb genomic windows enriched with derived alleles absent in Yoruba genomes (a sub-saharan modern human population with little or no Neanderthal introgression). Further refinement included segment delimitation by Altai Neanderthal-matching derived alleles, and removal of segments including derived modern human-specific alleles and with greater Denisovan than Altai Neanderthal match rate.

In our working dataset, which consisted in a cohort of 38,406 UK Biobank unrelated samples of European ancestry with available brain MRI scans and imputed genotypes, the introgressed segments present in at least 0.1% of individuals encompassed a total of 4,179,475 variants. Among these, we selected variants that are present in the UKBB only thanks to Neanderthal introgression. We first identified 352,663 variants where an allele matching the Altai Neanderthal had a frequency < 1% in Yoruba individuals. Then, similarly to previous studies^21^, we expanded this set by including variants in near-perfect linkage disequilibrium (LD) (R^2^ > 0.99 and distance < 200kb), thus identifying 422,195 Neanderthal Specific Variants (NSV). While most of these variants are Neanderthal-fixed derived (see **Figure 1A**), these steps ensure that Neanderthal polymorphisms, alleles specific to the introgressing Neanderthal, and indels poorly genotyped in the archaics are accounted for. When compared with pre-existing collections used to investigate Neanderthal phenotypic impact^17,19^ such as those from Browning et al.^4^ and Sankararaman et al.^6^, we include, respectively, 95.93% more and 60.56% more variants (**Figure 1B**, both sets were similarly LD-expanded in our sample set for a fair comparison). This increase is largely attributable to differences in sample size between earlier studies and the present work, as demonstrated by Morez Jacobs et al.^35^ by showing that the discovery rate of Neanderthal-introgressed variants continues to rise as thousands of genomes are added. However, part of this discrepancy may also reflect methodological differences in Neanderthal ancestry inference, such as reliance on Altai Neanderthal–specific alleles constrained to the strict 1000 Genomes mask^37^, as in Browning et al.^4^

**Figure 1.**
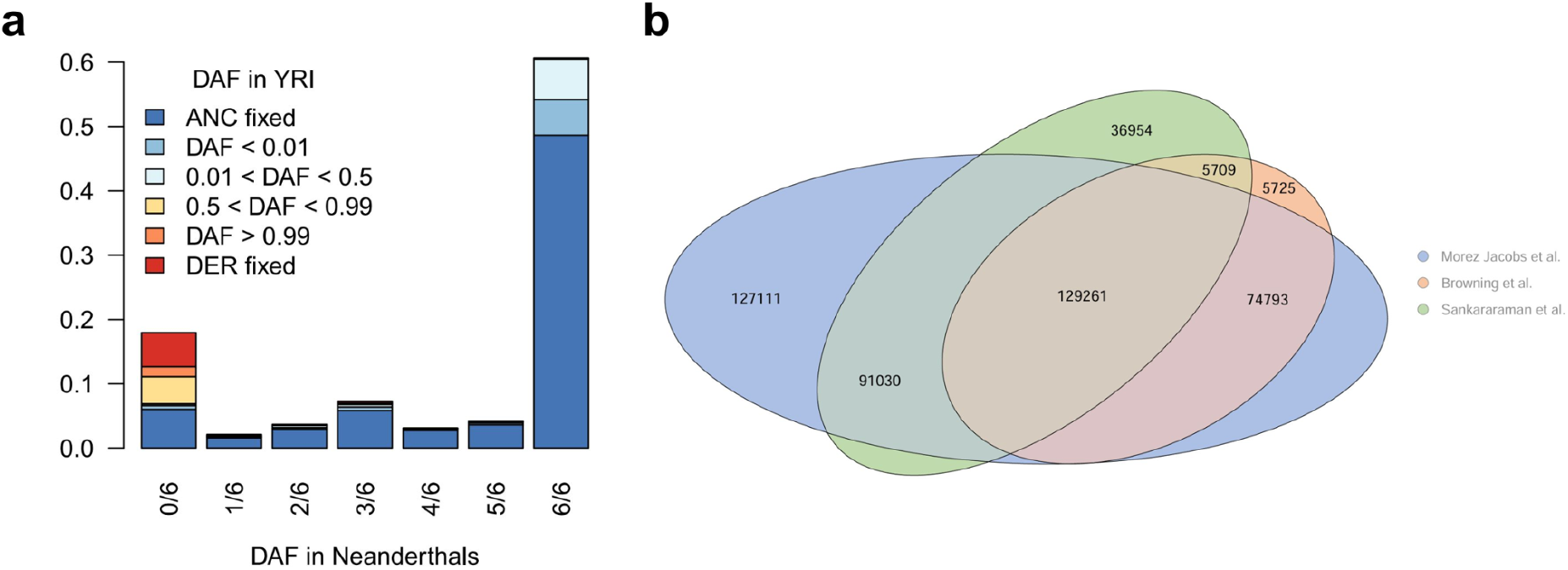
Neanderthal-specific variants. **a)** NSV derived allele distribution in a panel of three genotyped Neanderthals (x-axis) and in Yoruba individuals (YRI, color coded). Due to LD-expansion of NSV it is possible to have frequency of the Neanderthal allele > 0.01 in YRI, and even the same allele fixed in both Neanderthal and YRI, where the allele of the introgressing Neanderthal was different from the genotyped ones. **b)** Venn diagram of LD-expanded NSVs from three different sets: Browning et al.^4^, Sankararaman et al.^6^, and those identified in Morez Jacobs et al.^35^, adopted in this study.

### NSV contribution to brain morphology

Two recent studies converge on an overall depletion of heritability mediated by Neanderthal introgressed segments, which is naturally interpreted as post-admixture purifying selection against deleterious Neanderthal variations^17,19^. To ask whether this is also valid for MRI-derived brain morphology, we analyzed a set of 370 brain imaging-derived phenotypes (BIDPs) including regional cortical areas, volumes and thicknesses, subcortical volumes, cortical sulci measurements, and nerve tracts diffusivity quantifications (see **Table S1** for a full list). For each of these BIDPs, we performed a genome-wide association study (GWAS) in our cohort of 38,406 samples; details of this process are reported in Methods. Neuroimaging-derived phenotypes are often both phenotypically and genetically correlated^38–40^: see **Figure S1** for the phenotypic correlations we found among our set of BIDPs. We start, therefore, by investigating associations with this phenotypic domain as a whole, and we apportion the contribution to specific BIDPs in subsequent sections. To identify variants with a sizable effect in any of the analysed BIDPs, we combined the p-values resulting from GWAS using the aggregated Cauchy association test (ACAT)^41^ and selected suggestive associations at P_ACAT_<10^-5^. The ACAT procedure allows aggregating P-values derived from phenotypes with arbitrary correlation structure, and is thus appropriate for BIDPs.

In order to quantify the relative contribution of Neanderthal introgressed variants to specific phenotypes, it is crucial to control for the minor allele frequency (MAF) and total Linkage Disequilibrium (LD score^42^) joint distribution typical of NSVs: lower average MAF, higher LD score and absence of correlation between MAF and LD score^19^. To keep this into account, we asked whether the proportion of variants suggestively associated with any BIDP was different in specific MAF/LDscore bins for the following different categories: a) NSVs b) variants overlapping introgressed segments but not Neanderthal-specific (OVPint) c) variants in Neanderthal deserts as defined in Morez Jacobs et al.^35^ (NeaDes) d) remaining variants, mostly representing standing variation in common between the Neanderthal and modern human lineages (other). These proportions were obtained after LD clumping independently in each bin to avoid redundancy, and their error assessed with bootstrapping.

While this methodology is robust to NSV MAF and LD score joint properties, we could have alternatively applied the partitioned heritability approach adopted in Wei et al.^19^, which is inherently built to accommodate different genetic architectures. Nevertheless, it is also built for large sample sizes: the lower sample size available in UKBB for BIDPs compared to other traits (about 38 vs 350 thousands at the time of our investigation) leads to a large loss of power which makes the application of the same method less advisable (see **Supplementary Note 1**).

As shown in **Figure 2A**, MAF and LDscore increase the fraction of suggestively significant variants across all categories. Nonetheless, different variant categories show different significance fractions within the same MAF/LD score bin: at MAF 0.025-0.05 / LD score > 100, NSVs are depleted of significance (0 significant NSV) with respects to all other categories; at MAF > 0.25 NSV is the category with the highest fraction (33 significant NSVs in the high LD score bin). Neanderthal deserts also appear to be enriched in the latter frequency bin. If we fit significance at different thresholds with a logistic model including MAF and LD score (see Methods), NSVs do not appear to decrease significance compared to the NeaDes category for thresholds P<10^-5^ to P<10^-7^, as depicted in **Figure 2C**. Notably, Other and OVPint categories are always depleted of significance against the NeaDes baseline, and a strong depletion emerges for NSVs if we restrict to variants with MAF<0.25. This dual behaviour - whereby NSVs show depletion of significant variants at low MAF but escape depletion at higher MAF bins - is completely absent when we examine variant significance across all traits collected in the GWAS catalog^43^ (**Figure 2B**): here, the NSV significant fraction is nearly always lower than its Neanderthal desert counterpart, in agreement with the overall functional depletion of Neanderthal-introgressed variability. In a logistic model, here NSV indeed always has a significance-decreasing impact compared to NeaDes, after MAF and LD score contributions. Curiously enough, non-Neanderthal-specific variants overlapping introgressed segments have a GWAS catalog fraction similar to deserts, consistent with a purifying selection specific to NSV.

**Figure 2.**
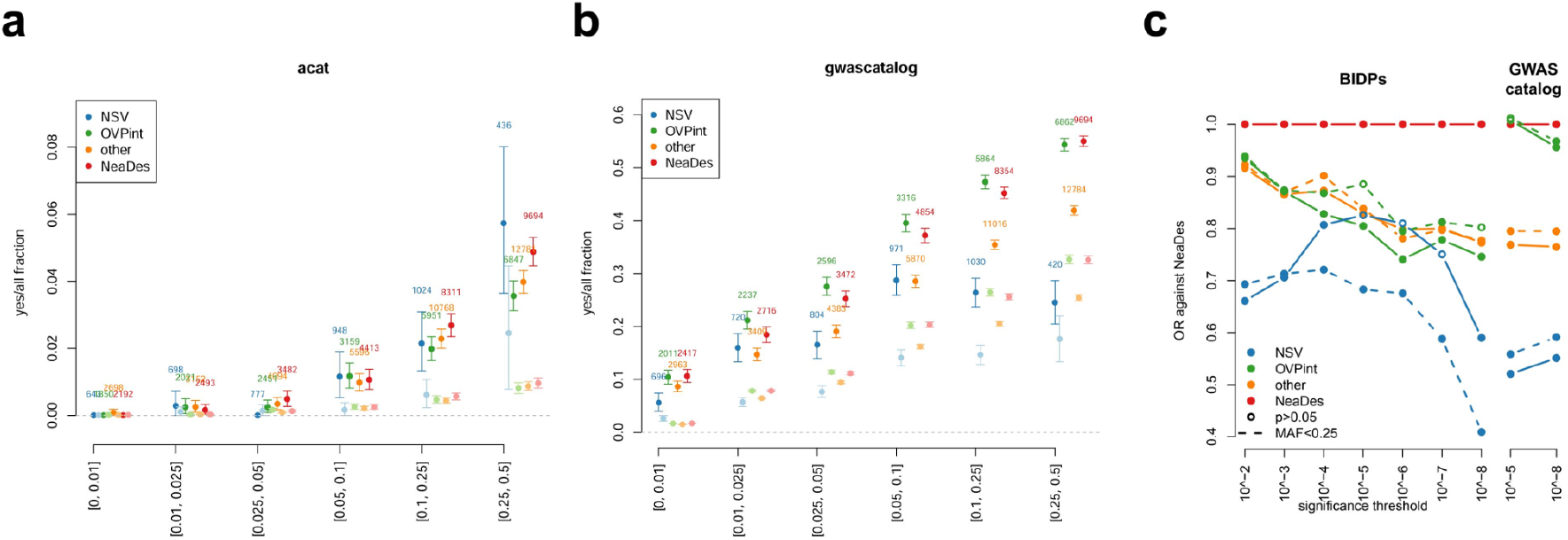
Fraction of NSV associated with BIDPs and with all traits in GWAS Catalog. **a)** variants suggestively associated with at least one BIDP (P_ACAT_<10^-5^) across four variant categories. Y axis: fraction of suggestively associated variants over the total number of variants in each category, the bars indicate 95% confidence intervals obtained through 10,000 bootstrapping runs. X axis: MAF bins. Dark and pastel points indicate bins of variants with high and low LD score respectively, with a cutoff set at LD score = 100. The numbers report the total number of variants in each category (i.e. the fraction denominator), for the high LD score bins, in each MAF bin. **b)** The same analysis showing the fraction of variants reported for any trait in the GWAS catalog. **c)** Odds Ratios (OR) for significance defined at different P-value thresholds, against Neanderthal desert category as baseline. ACAT P-values for BIDPs were used to determine significance on the left, minimum P-values reported in GWAS catalog on the right. OR were estimated by a logistic model including MAF and LD-derived covariates, see Methods for details. Dashed lines represent OR estimated discarding the top MAF bin, keeping variants with MAF<0.25.

These results suggest that while the majority of low-frequency NSVs are indeed devoid of suggestive contribution to BIDPs, a larger-than-expected fraction of NSVs with an impact on brain morphology appears to have escaped purifying selection (or even proven adaptive) and risen to medium-high frequencies, contrary to what is seen in most non-BIDP complex traits.

### NSVs with significant effects on brain morphology

We proceeded to identify the individual peaks of association significant at genome-wide level (P_ACAT_<5*10^-8^) that could be explained by Neanderthal introgression. Significant variants closer than 100 kb were merged together as they were likely tagging the same association. As shown in the Manhattan plot in **Figure 3**, we found 521 significant loci, out of which 28 (5.37%, **Table S2**) contained at least one significant NSV. Most of these signals replicated when testing the association between BIDPs and the Neanderthal dosage at each variant as inferred in Morez Jacobs et al.^35^ (see Methods). This is expected, as if the NSV indeed is the causal variant, the introgressed haplotype/trait association should closely mirror the GWAS signal. Indeed, at NSV the Neanderthal dosage matches the allelic status, so GWAS and Neanderthal dosage association signals should match as well.

**Figure 3.**
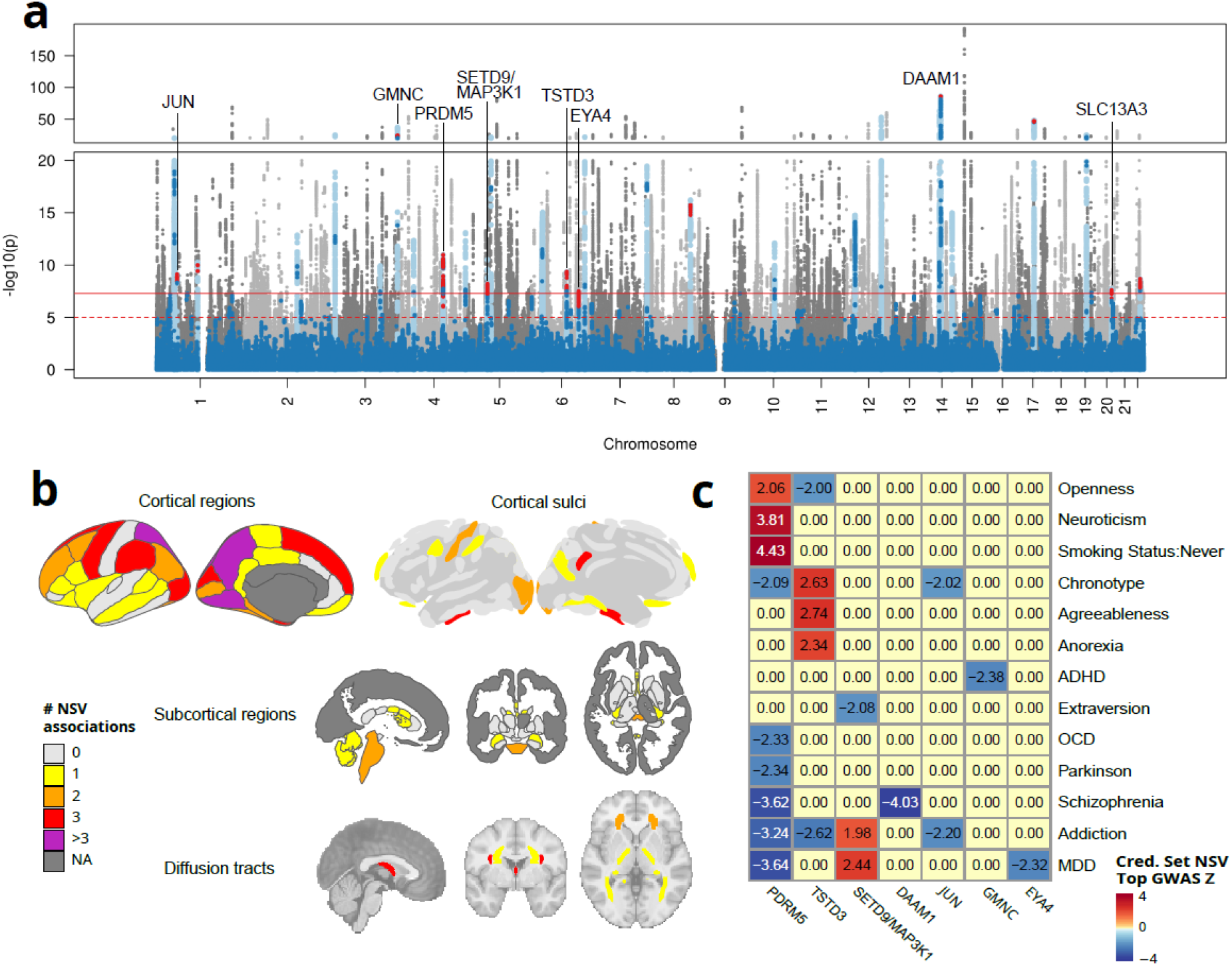
BIDP-associated NSV. **a)** Manhattan plot of the multi-trait ACAT analysis across all autosomes. Loci that contain at least one significant NSV are highlighted in light blue, NSVs are dark blue and NSV in credible sets are red. The eight loci with credible sets composed by a majority of NSVs are tagged with the name of a representative gene. Solid and dashed red lines represent the genome-wide significance (P_ACAT_<5*10^-8^) and suggestive significance (P_ACAT_<10^-5^) thresholds, respectively. **b)** Number of NSV-mediated associations for different brain regions (associations with different measurements are counted independently). **c)** Z-scores for associations between credible set NSVs and 17 mental health traits, oriented according to the Neanderthal allele. For each locus-trait pair, we report the Z-score of the most significant NSV-trait association within the credible set, provided it reached nominal significance, reporting 0 otherwise. ASD, bipolar disorder, alcohol use disorder, and conscientiousness showed no association with any NSV locus and are omitted from the figure.

To further corroborate the possibility that archaic alleles are indeed causative for some of these morphological associations we adopted two strategies. First, we used SuSiE^44^ to fine-map each of the 28 signals to the smallest credible set of likely causal variants: if the credible set did not overlap with the NSV, the causal variants are likely just in LD with a NSV. While 25 of these loci were significantly associated with multiple BIDPs, in 17 of them the majority of the associations colocalized with the top association, as is expected under pleiotropy among BIDPs (see **Table S3** for further details). For simplicity we therefore relied on the credible set for the top associated BIDP in each locus, with the caveat that we could lose some non-colocalizing

NSV-mediated associations in more complex loci. In eight loci, listed in **Table 1** and shown in **Figures 3, 4A, S2**, NSVs were the majority of the variants composing the credible set, confirming that Neanderthal introgressed variants are likely to be truly causative for the associated trait. Of these, seven loci were associated with multiple BIDPs and five had the majority of associations colocalizing with the top one.

**Table 1.** BIDP-associated NSVs. For each locus with a credible set (CS) composed by a majority of NSVs, the table reports a tagging gene, genomic coordinates (hg19), top NSV GWAS P_ACAT_, top P_ACAT_ from the Neanderthal dosage association analysis (NDA), associated traits CS size and percentage covered by NSVs, NSVs within a CS that either overlap an open chromatin region according to ATAC-seq or exceed the 99th percentile of the maximum absolute ΔSVM score distribution. Complete data are reported in **Table S4**.

| Tag gene | Chr | Start | End | Associated traits | Top NSV<br>$P_{\text{ACAT}}$ | Top NDA<br>$P_{\text{ACAT}}$ | CS<br>size | NSV<br>in<br>CS | CS<br>NSV<br>avg<br>MAF | CS NSV in<br>ATACseq<br>peak | CS NSV<br>with top<br>1% $\vert \Delta\text{SVM}\vert $ |
| --- | --- | --- | --- | --- | --- | --- | --- | --- | --- | --- | --- |
| JUN | 1 | 59217533 | 59432116 | posterior thalamic radiation FA +1 | 7.50E-10 | 6.43E-10 | 19 | 100% | 0.108 | rs3281,<br>rs61788403,<br>rs61788409<br>+2 | rs61788411 |
| DAAM1 | 14 | 59349137 | 60133892 | cuneus cortical volume +38 | 5.26E-87 | 1.20E-84 | 4 | 100% | 0.122 | rs74826997,<br>rs76341705 | NA |
| SLC13A3 | 20 | 45086396 | 45304763 | Superior longitudinal fasciculus MD +2 | 2.43E-08 | 3.42E-03 | 28 | 64% | 0.164 | rs2694876,<br>rs3091661<br>+10 | NA |
| GMNC | 3 | 190491314 | 190778438 | Ventricles and choroid plexus volume +35 | 2.56E-25 | 1.62E-24 | 42 | 81% | 0.096 | rs9818981 | rs9827488,<br>rs9828756 |
| PRDM5 | 4 | 121446341 | 121865788 | Paracentral cortical thickness +22 | 1.06E-11 | 2.71E-11 | 325 | 93% | 0.256 | rs11721741,<br>rs11733936<br>+14 | rs35013646,<br>rs3804169 |
| SETD9/M<br>AP3K1 | 5 | 55908915 | 56283743 | Brain stem volume +18 | 6.34E-09 | 6.66E-09 | 50 | 74% | 0.101 | rs1862620,<br>rs60985357<br>+3 | NA |
| TSTD3 | 6 | 99727313 | 100239801 | Cerebellum cortex volume | 4.80E-10 | 1.55E-08 | 53 | 98% | 0.215 | rs330872,<br>rs491178 +1 | rs12529031 |
| EYA4 | 6 | 133465734 | 133681377 | Pontine crossing tract L1 +11 | 3.12E-08 | 3.83E-08 | 51 | 75% | 0.222 | NA | rs9375942 |

**Figure 4.**
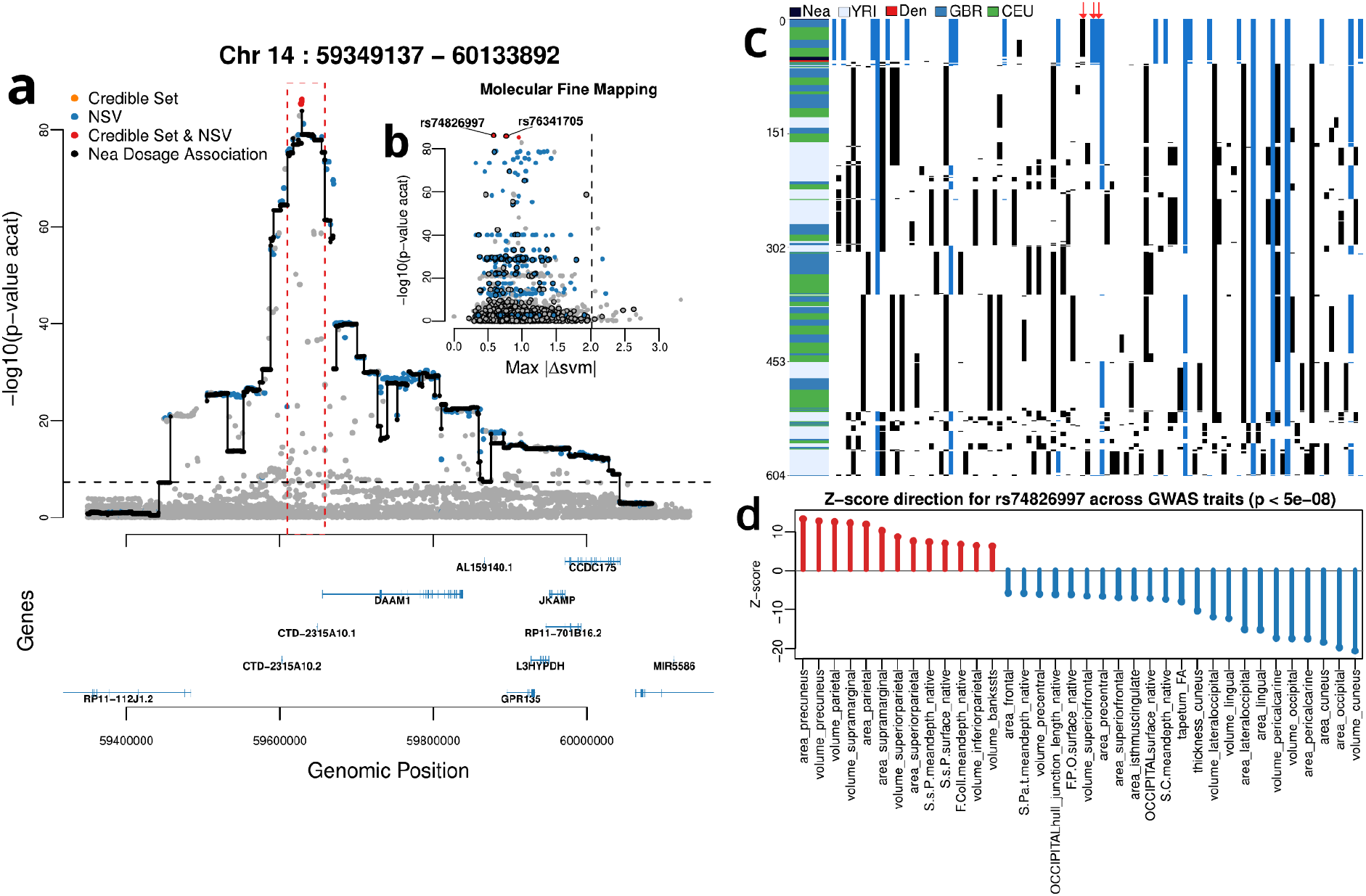
DAAM1 locus. **a)** Annotated locus zoom of the region chr14:59349137-60133892. Showing -log10(P_ACAT_) values for SNPs in the locus. Blue, orange, and red dots indicate,, respectively, variants that are NSVs, included in a credible set, and both; black dots connected by lines denote -log10(P_ACAT_) for Neanderthal dosage association. Gene annotations are reported in the bottom panel. **b)** Molecular fine mapping of the locus variants, using the same color coding and y-axis but highlighting variants overlapping ATAC-seq peak with a black contour, and reporting on the x axis the maximum absolute ΔSVM score. The dashed line represents the top percentile. Complete supporting data are reported in **Table S4. c)** Haplostrips^49^ visualisation of the introgressed haplotype at the DAAM1 locus, specifically the region delineated in red in **a)** (chr14:59610075-59658600). Each row represents a single phased haplotype, colour-labelled on the left by population (Nea: Neanderthal, Den: Denisovan, YRI: Yoruba, GBR:British, CEU: Utah Europeans, from 1000 Genomes panel). Each column represents a variant site, black cells indicate the derived allele, blue cells derived alleles in common with the Neanderthal haplotype and white cells ancestral alleles. Red arrows point to the positions of the credible set variants, rs2164950, rs74826997 and rs76341705 (14:59627434_AC_A is missing from the archaic VCF file adopted). **d)** Z-scores for genome-wide significant (p < 5e-08) associations of rs74826997 across BIDPs.

Secondly, we looked for a putative molecular mechanism for the credible set variants. We found no coding NSVs in any credible set, as expected since most of the GWAS signals fall within elements with regulatory activity^45^ in specific cellular and developmental contexts^46^ rather than altering protein sequences. We therefore asked whether NSV would fall within an open chromatin region identified by single-cell ATAC-seq on developmental and adult prefrontal cortex from Herring et al.^47^, or, alternatively, whether they would be predicted to alter chromatin accessibility by a gkmSVM model^48^ trained in the same cellular contexts. In all loci we have support for a plausible mechanism for their causality: in seven loci, at least one credible set NSV overlapped an open chromatin region in at least one cell type; in five loci, the maximum absolute ΔSVM^48^ score of one or more credible set NSVs fell within the top 1% of all SNPs tested. The variants involved are listed in **Table 1**, while we show in **Figures 4B, S3** the two molecular annotations for each locus.

One of our motivations is to investigate the parallelism between the Neanderthal impact on mental health and brain morphology. We thus asked whether the credible set identified showed any effect in 17 selected mental health GWASes (listed in **Table S5**). As visible in **Figure 3C** nominal effects were present, albeit sparse, in all loci except the one tagged by the gene *SLC13A3*. Numerous associations have been found instead for the credible set NSVs at the *PRDM5* locus which, among others, show protective effects against smoking and other substance addiction, major depressive disorder (MDD), schizophrenia, but a more neurotic and open personality.

Among all identified loci, a peak on chromosome 14 near the gene *DAAM1* stands out for its significance: we discuss it below. Other loci listed in **Table 1** are shown in **Figure S2**, and thoroughly discussed in **Supplementary Note 2**.

### Neanderthal-divergent variants at the DAAM1 locus

Chromosome 14 harbours the most significant association fine-mapped to a NSVs among the genome-wide significant loci: we tag it as *DAAM1*, its nearest gene (see Discussion for further details). Here, a large introgressed segment, spanning ∼585 kb (chr14:59457561-60042473) precisely overlaps the association signal (**Figure 4A**), including a shorter highly associated haplotype (chr14:59610075-59658600, 48.5kb, red dashed line in **Figure 4A**). The credible set lies within this haplotype, and encompasses four variants which are all NSVs: 14:59627434_AC_A, rs2164950, rs74826997 and rs76341705. Among these variants, all with a MAF=0.12 and fixed-ancestral in Yorubas, rs2164950, rs74826997 are fixed-derived in the three Neanderthals analyzed. rs74826997 and rs76341705 lie in a region of open chromatin in microglia and both excitatory and inhibitory neurons (see **Figure 4B**) so are the most likely causal candidates at this locus.

Analyzing the core haplotype pattern in **Figure 4C**, we can observe indeed a striking similarity to the Neanderthal consensus, and a complete absence of this haplotype in Yorubas, confirming its archaic origin. Moreover, this region harbors one of the strongest signatures of divergence between AMH and Neanderthal in the whole human genome^1^ (see **Figure S4**).

This locus is significantly associated with 39 BIDPs, including all occipital cortical areas and volumes, some parietal areas (superior parietal, precuneus, supramarginal), and a few dMRI tracts (tapetum and cingulum). While these include relevant regions in the Neanderthal-AMH braincase divergence such as the parietal lobe, the direction of the Neanderthal allele effect at the most likely causal variant (rs74826997, **Figure 4D**) is opposite to the standard of a reduced parietal region in archaic humans. Conversely, its protective effect against schizophrenia (**Figure 3C**) indicates this locus as one of the mediators of the previously identified inverse correlation between this condition and Neanderthal alleles^17^.

### Genome-wide directional effects of Neanderthal-derived variants

To assess whether, beyond individual loci, Neanderthal introgressed segments mediate a directional genome-wide effect, we implemented SLDP (Signed LD Profile) regression^50^. Starting from signed effect sizes estimated from 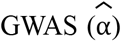, SLDP estimates *r*_*f*_, the correlation coefficient between these and a signed annotation, after accounting for LD. We applied this analysis to all BIDPs independently, using as annotation *Δp*_*Nea,YRI*_ : the effect allele frequency difference between a panel of three Neanderthal individuals (Altai, Vindija and Chagyrskaya) and a panel of Yoruba individuals from 1000 Genomes^51^. While naturally upweighting the effect of NSVs, this approach allows us to use all variants overlapping introgressed segments, thus accounting also for more subtle effects, and downweights the NSVs that are polymorphic within Neanderthals. As can be seen in **Figure 5A**, while we could not find any BIDP significant at FDR<0.05, some were significant at nominal level: in particular parietal (especially superior parietal), temporal fusiform and lateral orbitofrontal cortical areas; callosal and occipital sulcal lengths; external capsule and cerebral peduncle tracts diffusivity showed the most significant positive correlations with introgressed alleles. On the contrary, Neanderthal alleles were nominally negatively correlated with the volume of the inferior part of the lateral ventricle. Signal intensity is not a quantitative biological measure, we therefore avoid to interpret hypointense voxels in this paragraph.

**Figure 5.**
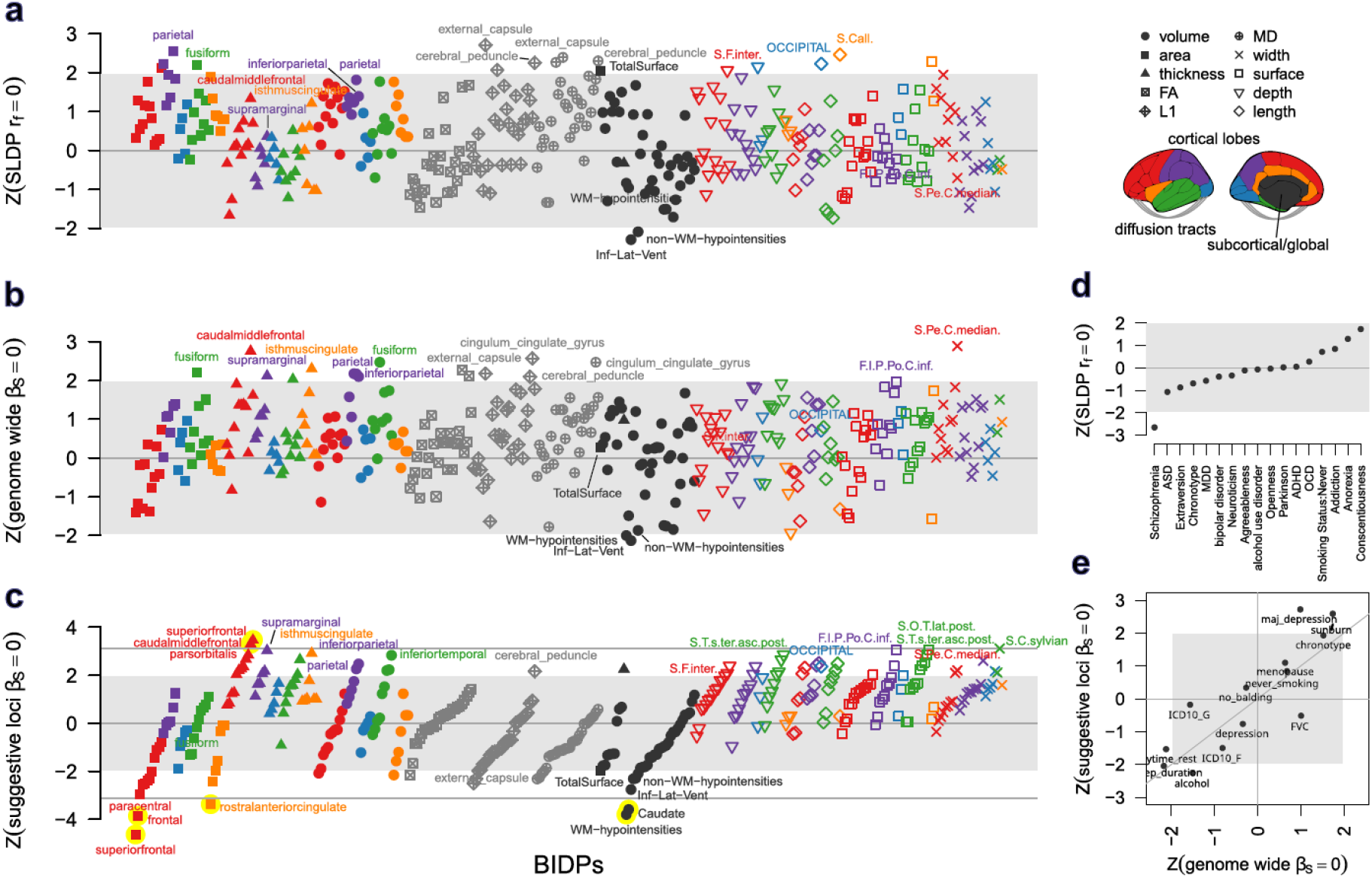
SLDP and Neanderthal burden correlations with BIDPs. **a)** SLDP Z statistic for the correlation (r_f_) between GWAS effect sizes and Δp_Altai,YRI_ annotation. **b)** Z statistic for Neanderthal burden correlation with BIDPs, computed on the whole genome. **c)** same as **b)** but computed on suggestive loci, i.e. 200kb windows centered on variants with P_ACAT_<10^-5^. In all these panels dots are shaped, colored and sorted according to BIDP categories and Z statistic from panel **c)**, labels are reported for top 5 BIDPs and nominally significant BIDPs recurring in more than one panel, dots shaded in yellow are significant at FDR<0.05. Grey shaded areas represent non-significant Z, while gray lines represent thresholds for FDR<0.05. **d)** SLDP Z statistics for the correlation between behavioural/neuropsychiatric GWAS effect sizes and Δp_Altai,YRI_ annotation. **e)** Z statistic for Neanderthal burden correlation with behavioral traits, health outcomes, and reference traits previously indicated as affected by Neanderthal introgression. Complete data are reported in **Tables S7-8**.

SLDP estimates of the SNP-based heritability of these traits are consistent with previous ones^33^, but the heritability attributed to *Δp*_*Nea,YRI*_ annotation is always < 5*10^-6^ (and often negative), well below the 1.2*10^-3^ estimate of heritability mediated by NSV from Wei et al.^19^ for a diverse panel of traits (see **Figure S5**). This suggests that SLDP struggles to capture any effect attributed to Neanderthal variability as quantified by *Δp*_*Nea,YRI*_, either due to a radical depletion in the heritability it mediates (which does not seem to be the case at least for a sizable set of high-frequency variants) or due to the combined limitations presented by the small fraction of annotated variants and a lower GWAS sample size for BIDPs than other traits. We reasoned that by moving from a summary-based approach to an individual analysis we could overcome the limitations due to the large uncertainties in GWAS summary statistics.

Akin to a similar approach already adopted by Gregory et al.^32^, we thus quantified each individual’s Neanderthal genomic burden and tested whether this score could predict any BIDP in a model including the same covariates as the GWAS study. As Neanderthal burden we adopted an individual’s count of alleles which are present in at least a Neanderthal and at frequency p<0.01 in Yorubas. This approach is similar to an enhanced D-statistic^52^, as it only factors in sites where an outgroup (Yorubas) does not carry the introgressed allele, but also leverages shared ancestral variation between the donor population and the recipient population, similarly to D+ statistic^53^. As shown in **Figure 5B**, no associations were significant at FDR<0.05 although several showed nominal significance. Nevertheless, even under a polygenic model, most of the genetic basis of BIDPs is likely concentrated in a small portion of the genome; it is therefore conceivable that a large fraction of the variance in Neanderthal burden is completely irrelevant to our phenotypic domain. As in other analyses^54,55^, we therefore resorted to limiting the burden calculation to a set of trait-associated genomic regions, selecting 200kb windows centered on loci suggestively associated to at least one BIDP (P_ACAT_<10^-5^). Genome-wide burden results were correlated with SLDP results (r=0.41, **Figure 5D**): this correlation faded away with stricter selection of BIDP-relevant loci. Nevertheless, for different parameter choices in selecting BIDP-associated regions the results remained qualitatively similar (r=0.64-93).

As shown in **Figure 5C**, the local Neanderthal burden computed on these variants was associated with a significant decrease in four biologically relevant BIDPs after multiple testing correction: superior frontal cortical area, frontal lobe area, rostral anterior cingulate area, volume of the caudate nucleus. Neanderthal burden also has a positive impact on the thickness of the same frontal cortical regions, suggesting a divergent arrangement in the grey matter of the frontal lobe. The same tendency is visible across the whole frontal and cingulate cortex, but not elsewhere, with parietal and temporal fusiform regions being consistently thicker and wider. Some sulcal measurements were positively associated with Neanderthal burden at FDR<0.1: the intraparietal section of the postcentral sulcus and two other temporal sulci internal surfaces, and the lateral end of the central sulcus width.

While the Neanderthal impact on mental health has been previously investigated^17,18,21^, we wanted to assess whether with the methods we adopted we can identify these signals. First we applied SLDP on a set of 17 behavioral and neuropsychiatric GWAS meta-analyses, finding the protective effect against schizophrenia (P=0.0079) previously reported by McArthur et al.^17^, but failing to replicate their signal on anorexia (see **Figure 5D**). Then, we regressed on the individual Neanderthal genomic burden a selected set of traits with a sufficient effective sample size in our UKBB cohort (**Table S6**). The most significant signal, specifically emerging when computing the Neanderthal burden on BIDP-relevant genomic regions, was a positive association with MDD (P=0.0065, see **Figure 5E**): this suggests a possible health consequence of the same introgressed variants implicated in the morphological changes we investigated.

## Discussion

Our analysis of UK Biobank neuroimaging data reveals that Neanderthal-derived variants exert a complex influence on present-day brain morphology. Although low-frequency NSVs show the expected depletion of significant associations - consistent with strong post-admixture purifying selection - this pattern is lost at higher frequencies. Indeed, 28 loci contain genome-wide significant NSVs influencing diverse imaging-derived phenotypes. This indicates that while most introgressed alleles affecting brain structure were removed, a subset either escaped purifying selection by conferring effects compatible with the modern human developmental context or were actively retained through positive selection, potentially because they offered adaptive advantages in novel post-admixture environments.

Fine-mapping strengthens this interpretation. In 8 loci, NSVs compose the majority of credible sets of likely causal variants, and in all but one cases at least one NSV overlaps an open chromatin region in developing or adult cortical cell types. Several NSVs rank among the top predicted chromatin-altering variants, suggesting that introgressed alleles can influence brain morphology through regulatory mechanisms.

The DAAM1 locus on chromosome 14 exemplifies this pattern. This locus has been previously recognized as associated with occipital and parietal cortical measurements^56,57^: here we demonstrate that this association is mediated by a ∼48 kb Neanderthal introgressed haplotype. This segment contains four credible-set NSVs and is associated with 39 traits spanning occipital and parietal cortices, including the precuneus and superior parietal lobule. These regions are among the most divergent between Neanderthals and modern humans, yet the direction of effect at this locus does not simply recapitulate the smaller precuneus and larger occipital lobe expected from Neanderthal cranial morphology. Instead, it suggests opposite effects on cortical patterning. According to GTEx^58^, the introgressed variants are eQTLs for DAAM1 and two downstream genes (L3HYPDH, JKAMP). In addition to a strong AMH/Neanderthal divergence at this locus^1^, a regulatory variant targeting DAAM1 documented to have emerged around 40 kya and subsequently risen above 90% frequency^59^ further supports its recent expression reprogramming. DAAM1 known roles in cytoskeletal dynamics and morphogenesis, Wnt–RhoA/ROCK signaling^60^, and the existence of a conserved neuronal microexon linked to synaptic plasticity and memory formation^61^ provide a convincing mechanistic route. Associations of these variants with diffusivity along tapetum and cingulum tracts further imply effects on long-range connectivity, while a protective effect against schizophrenia underlines this locus consequences on mental health.

Beyond DAAM1, several additional loci illustrate how introgressed regulatory variants may have influenced diverse neurodevelopmental pathways. On chromosome 4, a large credible set composed at 93% by NSVs with an extensive open chromatin overlap and a large potential to alter chromatin accessibility in cortical neurons acts as eQTLs and sQTLs for PRDM5 (GTEx).

These variants are associated with widespread frontoparietal cortical thinning, consistent with PRDM5’s evolutionarily conserved role in Wnt/β-catenin regulation during early development^62^.This locus also shows broad impact on MDD and other addictive and personality disorders. Notably PRDM5 and JUN, another gene overlapping a NSV-mediated association with the posterior thalamic radiation tract, have been identified in a study of archaic cis-regulated transcription factors as the only two factors with predicted target genes enriched in Neandertal deserts^63^.

Another notable example is the GMNC locus on chromosome 3, which shows one of the broadest effects, spanning from cortical and sulcal measurements, diffusion across several tracts such as the fornix, ventricular, choroid plexus and other subcortical volumes, in line with GMNC’s key role in multiciliated ependymal cell differentiation and CSF homeostasis^64^. Here two NSV on a secondary credible set are very likely to alter chromatin accessibility in microglia, oligodendrocytes, and MGE interneurons. The last credible set with a clear gene candidate contains eQTLs for SLC13A3, a gene predominantly expressed in astrocytes and previously associated with leukoencephalopathy with episodic neurological crises^65^. Given its role in white-matter integrity, it is notable that the locus associates with diffusion-MRI measures, including superior corona radiata MD and superior longitudinal fasciculus FA/MD. Remaining NSV-mediated credible sets contain eQTLs for less brain-characterized genes such as SETD9, EYA4, TSTD3 and others. However, these loci illustrate how surviving Neanderthal alleles map onto diverse BIDPs and underlying developmental systems, from cortical patterning to subcortical organisation and long-range white matter architecture.

When testing for genome-wide directional effects, we observed several nominal trends, but few were significant after multiple testing. Specific regions of the frontal cortex exhibited divergent thickness and area directions, temporal fusiform and parietal cortices were often among the top ranking BIDPs. Several sulcal measurements and diffusion tracts also were nominally associated, but none showed consistency across methods except for diffusivity along the cerebral peduncle. Notably, one of our strongest signals involves the caudate nucleus, hinting that evolutionary divergence between Neanderthals and modern humans may involve internal anatomical features invisible to endocasts. While the most significant signals appeared when restricting the Neanderthal burden analysis to BIDP-associated loci, this represents a drastic simplification and may leave out BIDPs with a highly polygenic architecture. Conversely, the fact that MDD associates with Neanderthal burden only under this restriction highlights a previously unrecognized pathway linking Neanderthal-derived variants to MDD through brain-morphology intermediates, warranting further investigation with e.g. Mendelian randomization approaches.

Overall, the depletion of low-frequency NSVs with measurable effects, together with the absence of genome-wide patterns matching the globularization process, supports a model in which most Neanderthal variants affecting brain morphology were deleterious in the AMH background. This is consistent with the known enrichment of neurodevelopmental genes in Neanderthal deserts^15^. However, high frequency surviving alleles, exemplified by the DAAM1 locus, illuminate how regulatory variation inherited from Neanderthals can still shape cortical organization and connectivity in subtle but biologically meaningful ways, with consequences on neuropsychiatric conditions such as schizophrenia and MDD. These findings underscore the value of introgressed variation as a window into the evolutionary forces that shaped the modern human brain.

## Materials and Methods

### Identification of introgressed segments

Introgressed segments were identified as described in Morez Jacobs et al.^35^. In brief, 45,532 donors with available imputed genome (Field ID 22828) and brain MRI-derived phenotypes as of October 2023 (ASEG whole brain volume, Field ID 26514, used as representative trait for filtering) were selected from UKBB. SSTAR^36^ was used to estimate S* score for 50-kb windows with a 10-kb step size, using the Yoruba genomes from 1000 Genomes panel^51^ (1000G) as reference and the UKBB genomes as target. S* score identifies divergent haplotypes in a target population that carry derived variants in strong linkage disequilibrium (LD) while being absent in the reference population^5,13,66^. Segments where the observed S* score fell within the top 99^th^ percentile of a null distribution generated under a neutral model without introgression were retained.

Putative Neanderthal segments were delimited by the first and last Neanderthal variants, i.e. variants present in the Altai Neanderthal genome and with a derived allele frequency (DAF) < 1% in the Yoruba genomes, segments without these features were removed. The refinement further included: a) merging overlapping segments; b) deleting any segment with a modern human variant (i.e. variants with a DAF > 1% in the Yoruba genomes and absent in the Altai Neanderthal genome); c) removal of segments with a match rate to Denisova 0.01 greater than with the Altai Neanderthal. Further information is available in Morez Jacobs et al.^67^.

### Definition of NSV and other variant categories

We first filtered the set of introgressed variants identified by SSTAR with frequency of introgression > 0.001. We then defined our set of Neanderthal Specific Variants (NSV) as introgressed variants with frequency of one of the two alleles in Altai Neanderthal > 0 and in African Yoruba <0.01. We expanded this set by LD using plink (with the flags --r2 --ld-window-kb 200 --ld-window-r2 0.99). All the remaining introgressed variants fall into the definition of OVPint, i.e. overlapping introgressed segments, but not Neanderthal-specific. Considering the non-introgressed variants, we generated a subset termed Neanderthal deserts (NeaDes), based on the definition of “expanded gaps” provided in Morez Jacobs et al.^35^ and covering 35.1% of the genome. We labelled the whole rest of non-introgressed variants as “others”.

### Genetic data for GWAS and subsequent analyses

Among the 45,532 UKBB samples with brain MRI-derived phenotypes described above, we selected samples identified as British with West-European ancestry (field 22006, code 1), not considered outliers for missingness and heterozygosity (field 22027), pruned to remove kinship coefficient ≥ 0.04419 (third or lower degree relatives). This resulted in a sample size of 38,406, with 16,159,788 variants retained for having MAF in UKBB >0.001 and imputation INFO score > 0.8.

In addition, to estimate allele frequencies in Neanderthals and Yoruba we used the variant calls for three high-coverage Neanderthal genomes from Altai^2^, Vindija^68^ and Chagyrskaya^69^; and 108 samples from 1000G phase 3 tagged as YRI.

### Phenotypic data

For each of these samples we analyzed 370 brain MRI-derived phenotypes listed in **Table S1** and including:

1. areas, volumes and thicknesses (33 measures each) of cortical regions segmented according the Desikan-Killiany atlas^70^; measures were averaged across left and right hemispheres;
2. for each lobe (frontal, parietal, occipital, temporal, cingulate, insular), we computed the total area, volume and average thickness (weighted by ROI area) yielding 18 lobar phenotypes;
3. 30 subcortical regions volumes generated via FreeSurfer automatic segmentation (aseg), again averaged across hemispheres when bilateral.
4. 11 global brain measures including: intracranial volume (ICV), Desikan atlas TotalSurface and GlobalMeanThickness descriptors, several aseg descriptors such as BrainSeg, Cortex, TotalGray, etc
5. three diffusion MRI measurements (fractional anisotropy, mean diffusivity and L1 axis) computed by Tract-Based Spatial Statistics for 27 diffusion tracts which reflect structural connectivity between pairs of brain regions;
6. 131 sulcal measurements obtained by averaging left and right hemisphere measurements of reliable sulci according to Pizzagalli et al.^71^ as described below.

While other measurements were already generated based on UKBB MRI images^33,38^ and made available as pre-computed phenotypes, sulcal measurements were computed according to the ENIGMA-Sulci protocol (https://github.com/ENIGMA-git), a standardized framework described in previous studies^40,71,72^ . This protocol relies on the BrainVISA 5.1.2 software environment^73^, which integrates several processing tools, including the Morphologist pipeline^74–76^. Starting from structural images provided by the UK Biobank, processed to replicate the UKBB MRI pipeline, FreeSurfer-derived segmentation was integrated in Morphologist to improve sulcal extraction, identification, and morphometric quantification. This fully automated procedure segmented and labeled 123 cerebral sulci (62 in the left hemisphere and 61 in the right hemisphere) according to the BrainVISA atlas. For each identified sulcus, morphological descriptors were computed in native (individual-subject) space. Sulcal length was estimated by counting the voxels located at the junction between the sulcus and the brain hull. Mean depth was defined as the average geodesic distance from the sulcal fundus to the brain hull. Surface area corresponded to the total area of the sulcal mesh. Mean width was calculated as the ratio between the volume of cerebrospinal fluid enclosed within the sulcus and its surface area^71,74,77^. Of the possible 492 measurements, we selected the reliable ones identified in Pizzagalli et al.^71^ and computed inter-hemisphere averages, thus obtaining 131 phenotypes. All sulcal labels are reported using BrainVISA nomenclature and described according to the “Atlas of the cerebral sulci” by Michio Ono et al.^78^.

### GWAS

Variants were further filtered for the GWAS based on genotype missingness (> 5% removed), minor allele frequency (< 0.001) and Hardy-Weinberg equilibrium (<10^-7^) thresholds using PLINK2^79^, resulting in a set of 12,156,331 tested variants. For the association we adopted REGENIE^80^: REGENIE performs GWAS chromosome-by-chromosome including polygenic predictions from other chromosomes for robust correction of population stratification. Variants were filtered prior to polygenic model generation to retain only well-imputed and high-quality markers. We restricted the set of variants with imputation INFO scores > 0.90 and applied linkage disequilibrium (LD) pruning (2000 Kb window with 1 SNPs as spacing, r^2^ > 0.5) to obtain a subset of variants in approximate LD equilibrium. This reduced the set of variants for REGENIE step 1 to 781,093 SNPs. The polygenic predictors estimated in step 1 were then used in REGENIE step 2 to perform GWAS on all variants.

The confound set included variables for head size, brain position within the scanner, age, sex and imaging site^38^, as well as 20 population genetic principal components (see **Table S9** in Supplementary Tables). Principal components were computed on all variants after LD pruning (1000 Kb window, r^2^ > 0.1) using the *pca* tool from PLINK2.The non-site, non-genetic confounders were processed and expanded in a manner similar to that carried out by Alfaro-Almagro et al.^81^. In brief, an initial set of 12 confounders was expanded to 34 through a series of site-interaction processing steps. Quantitative confounds were de-medianed and normalised globally, then replicated per site so that each copy retained only values for subjects scanned at the corresponding site. Outliers and missing values were replaced by the site’s median before normalisation. Categorical confounds were handled by binarisation into indicator variables and replicated by site where more than 1 unique value was observed, before being de-medianed by site. Overall, this resulted in a set of 56 confounding variables.

A Neanderthal dosage association analysis was performed to identify the phenotypic effects of Neanderthal-introgressed haplotypes in European individuals with detectable Neanderthal burden, following the same analytical framework as the primary GWAS, with three key modifications. First, we started from a variant file obtained from Morez Jacobs et al^35^. which reported the ancestry status instead of the allelic status, with 1 representing alleles annotated to introgressed segments and 0 elsewhere. Second, the Hardy-Weinberg equilibrium filter was omitted, as introgressed Neanderthal haplotypes violate multiple assumptions of HWE, including the absence of gene flow, random mating and negligible genetic drift, and would therefore be systematically flagged and removed by this filter. Third, we used the set of high-quality, LD-pruned SNPs derived from the primary GWAS for the regression model in

REGENIE step 1, as the number of variants within Neanderthal-introgressed haplotypes alone was insufficient to fit the whole-genome regression model. Using the full set of SNPs from the full discovery cohort also ensures a solid genetic genetic relationship matrix and polygenic background estimate.

For both associations, P-values across traits were combined with the aggregated Cauchy association test (ACAT)^41^ and the resulting P-value (P_ACAT_) was used to identify significant associations in at least one trait.

### Intersection of NSV with BIDP-associated and GWAS catalog variants

We separated variants in our dataset in the four different categories (NSV, OVPint, NeaDes and others). Since we need the variants in each subset to be independent and to preserve the significant variants we performed LD clumping (r^2^ cutoff = 0.2, over a window of 250 kb), separately for each category. Then we divided them into 12 bins: 6 MAF bins (breaks: 0,0.01, 0.025, 0.05, 0.1, 0.25, 0.5) and 2 LDscore bins (splitting at LDscore 100). For each bin, we then computed the fraction of NSV, OVPint, other and NeaDes variants that resulted in significant associations from the ACAT analysis (P_ACAT_<10^-5^), over the total number of variants of the same category in that bin. We obtained 95% confidence intervals on the fraction with 10,000 bootstrap rounds.

We performed the same analysis, this time selecting as significant variants annotated in the GWAS catalog database (a comprehensive collection of human Genome-wide association studies). Finally, to estimate the effect of different categories relative to deserts we ran the following logistic model : significance status ∼ MAFbin + MAF + MAFbin:MAF + LDscorebin + LDscore + LDscorebin:LDscore + variant category.

### Statistical and molecular Fine-mapping

Fine-mapping was conducted using the Sum of Single Effects model implemented in the susieR R package^82^. For each locus, fine-mapping was performed separately for all MRI traits showing genome-wide significant associations within the region, using GWAS summary statistics and linkage disequilibrium (LD) estimated from the study sample. The SuSiE model was fitted using the *susie_rss* function, allowing up to 10 causal variants per locus (L = 10), with a sample size parameter of n = 38,000 and a maximum of 300 iterations. Credible sets where NSVs represented the majority of variants were considered Neanderthal-driven. This rationale is analogous to the one used in Wei et al.^19^, although we considered credible sets at the same locus independently.

To assess whether association signals across traits reflected shared or distinct causal variants, we performed pairwise colocalization analyses. Within each locus, colocalization was evaluated between each significant MRI trait and the top trait, defined as the phenotype harboring the SNP with the lowest GWAS p-value in the region. Colocalization was performed using *coloc*.*susie* implemented in coloc R package^83^ applied to SuSiE fine-mapping results, which allows for multiple causal variants per locus. Default prior probabilities were used. The colocalization hypothesis was considered as supported when PP.H4 had the highest posterior probability.

To identify putatively regulatory regions we adopted single cell ATAC-seq called peaks from Herring et al.^47^. These include 9 brain cell types (astrocytes, microglia, upper-layer excitatory neurons L2/3, deep-layer excitatory neurons L5/6, CGE-derived interneurons, L4 excitatory neurons, MGE-derived interneurons, oligodendrocytes and vascular cells) and developmental stages from gestational week 22 to adulthood.

The same peaks were used to train gkm-SVM^48^ models, which were then used to predict the effect of each single nucleotide variant on the chromatin accessibility in each cell type using ΔSVM scores. Each model was trained on 5,000 open chromatin regions randomly drawn from the initial dataset, with each sequence limited to a maximum length of 8,000 base pairs, all default parameters were used. In total, 84,145 SNPs were analyzed, derived from 28 loci that each contained at least one significant NSV. To identify the set of variants with a putative effect we computed the maximum absolute ΔSVM across cell types and adopted as global threshold the 99th percentile (ΔSVM=2.0179) across all SNPs tested. Variants exceeding this threshold or overlapping ATAC-seq peaks were flagged as candidates of functional relevance.

### Directional analysis based on Signed LD Profile regression

SLDP^50^ regression estimates the correlation between signed summary statistics from a GWAS and a signed annotation after computing the annotation LD profile, i.e. the marginal annotation signals carried by LD. SLDP estimates parameters in this model:

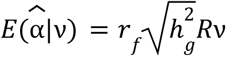

where 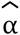 is the vector of marginal correlations between SNP alleles and a trait, *v* is the signed functional annotation, *r* _*f*_ is the functional correlation mentioned above, 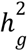 is the SNP heritability of the trait of interest, and R is the LD matrix. The model is tested against a signed background model, corresponding to the directional effects of minor alleles divided in MAF bins, necessary to address confounding effects due to genome-wide directional bias of minor alleles due to selection.

We adopted as *v* the difference in frequency between a panel of three Neanderthals and 108 YRI individuals from 1000G, signed so that the annotation is positive when the GWAS effect allele corresponds to the allele most frequent in Neanderthals and negative otherwise (*Δp*_*Nea,YRI*_ *=* p_Nea_ - p_YRI_). This scoring was restricted to the set of variants overlapping introgressed segments, while a score of 0 was assigned to all the non-introgressed variants. This scoring results in a positive *r*_*f*_ when the Neanderthal alleles are correlated with a trait increase, and a negative *r*_*f*_ when the Neanderthal alleles are correlated with a trait decrease.

In order to reduce the computational load of our analysis, we performed genome-wide LD pruning with a very relaxed cutoff (r^2^>0.8), and in order to discard alleles with a large error in the effect size estimate we restricted our analysis to SNPs with MAF > 0.01.

Neanderthal burden scores were computed using plink -score function relying on NSV where the allele with frequency p<0.01 in Yorubas had p>0 in Neanderthals, removing NSVs not satisfying this condition. The scoring weight was 1 for alternative alleles matching the Neanderthal allele and -1 for alternative alleles matching the Yoruba allele. This is equivalent to the NeanderScore used in Gregory et al.^32^ and to an Enhanced D-Statistic^52^ that also includes ancestral Neanderthal alleles in addition to derived Neanderthal alleles. This addition accounts for re-introduction of ancestral variants lost due to fixation of derived alleles in AMH, and is expected to increase the precision in estimating individual neanderthal burden^53^. These scores were used as a predictor of the trait in a linear model including all covariates used in the GWAS for BIDPs, and including sex, age, age^2^, sex:age, sex:age^2^, site and the first 20 genetic PCs for non-BIDP traits.

## Acknowledgements

This research was funded by the European Union - Next Generation EU, Mission 4 Component 1, CUP D53D23016690001. This research has been conducted using the UK Biobank Resource under application number 86275.

## Data Availability

The data adopted in this research were either publicly available or available under controlled access by UK Biobank Resource. The summary data generated not already accessible in supplementary materials (individual BIDP GWAS and Neanderthal Dosage Associations) will be available upon acceptance. The individual-based data generated will be made available through the UK Biobank data return program.

## Code Availability

The analyses here performed used publicly available tools cited throughout the manuscript. All the pipelines used to generate the data and figures will be uploaded at the address https://github.com/dmarnett/neanderthalinourbrain upon acceptance.

