## Supplementary Material for "Effects of introgressed Neanderthal alleles on present-day brain morphology"

#### Supplementary Figures

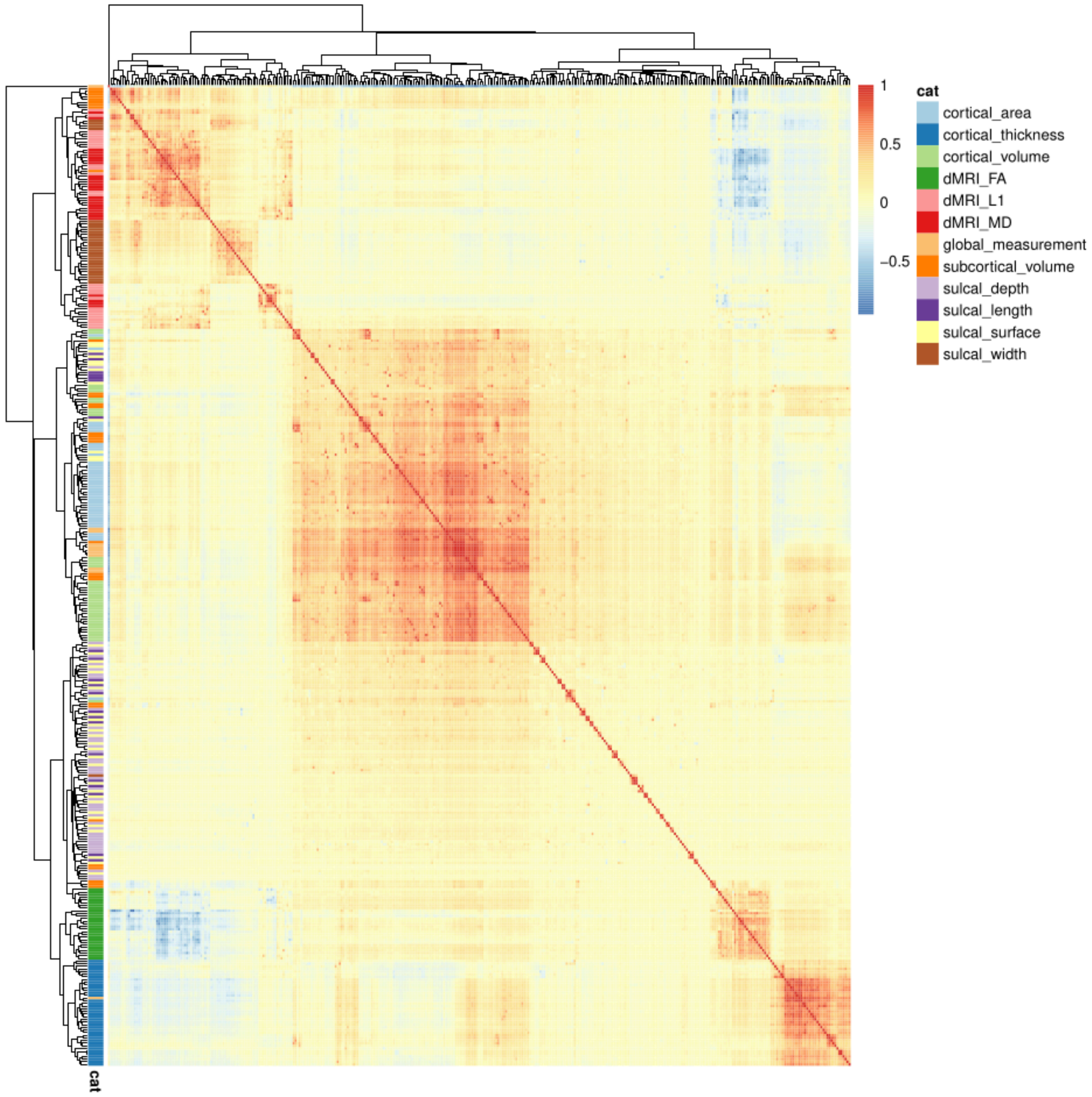


***Figure S1. Phenotypic pairwise correlations between BIDPs.*** *Pairwise correlations computed on the UKBB dataset adopted in this paper.*

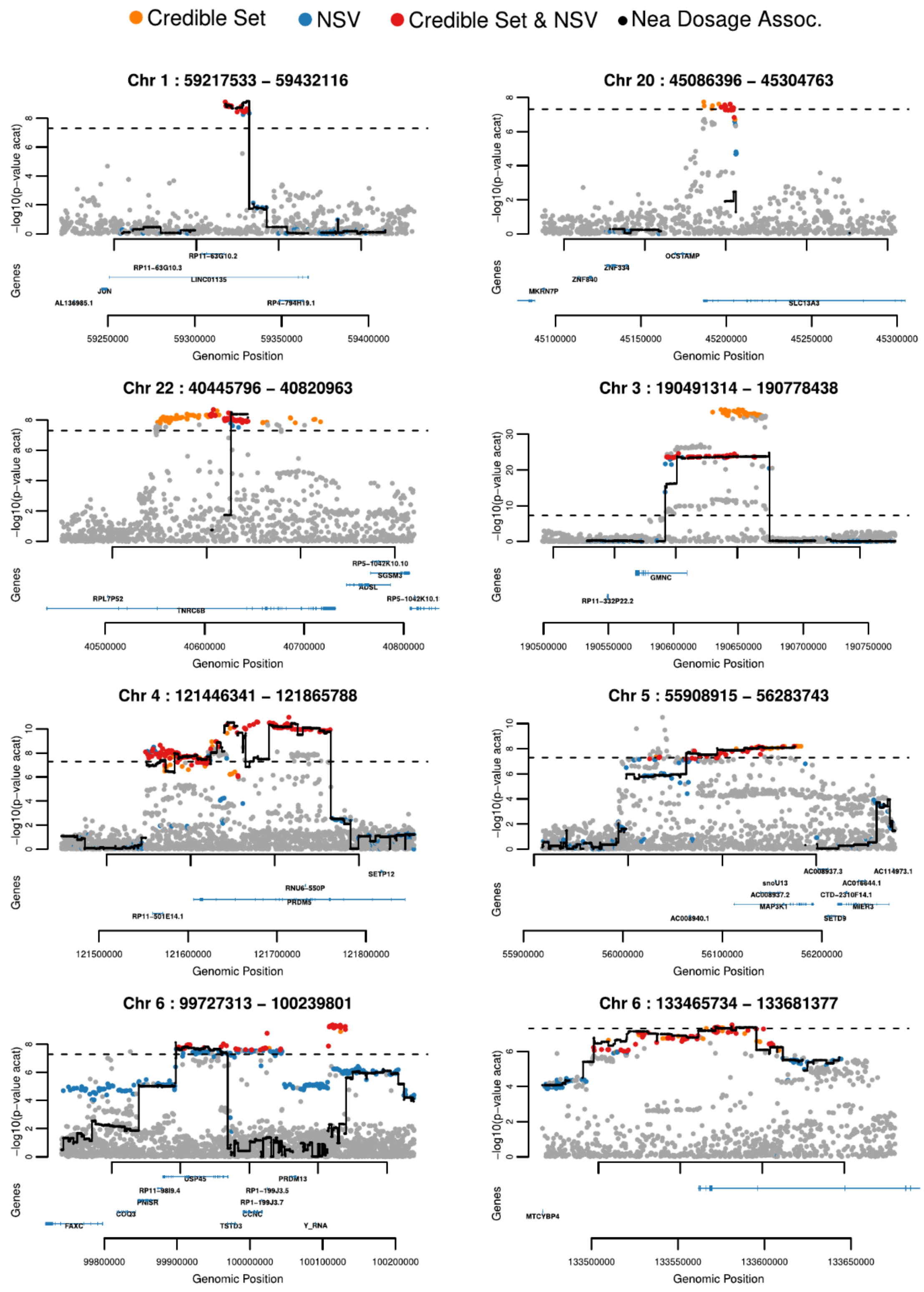


***Fig S2. Locus zoom plots of Neanderthal associated regions bearing NSVs in a SuSiE credible set.*** *Each panel displays an annotated locus zoom plot for seven loci bearing more than 50 % of NSVs within a SuSiE credible set. In addition we include the locus 22:40,445,796-40,820,963 whose credible set is composed for the 41% by NSVs as a suggestive case. Blue, orange, and red dots indicate NSV, variants in a credible set, and both, respectively; black dots connected by lines denote -log10($P_{ACAT}$) for Neanderthal dosage association. Gene annotations are reported in the bottom panel. (excluding 22:40,445,796-40,820,963, bearing more than 40 % of NSVs).*

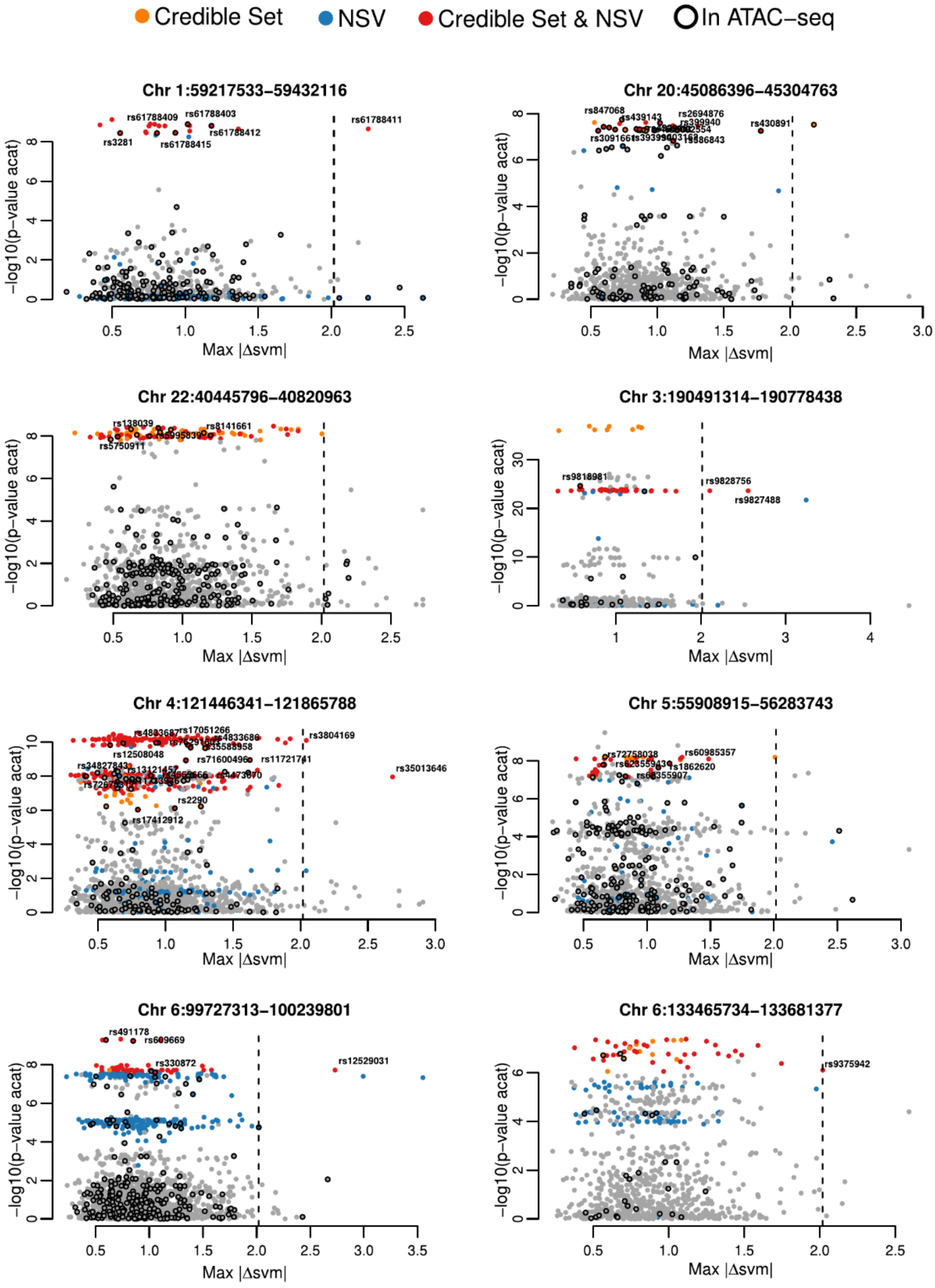


***Fig S3. Fine-mapping of Neanderthal associated regions.*** *Each panel displays variants within a genomic locus plotted by their maximum |Δsvm| score (Max |Δsvm|) against their association significance ($-\log_{10} P_{ACAT}$). Points are coloured according to variant category: orange, credible set (CS) variants; blue, Neanderthal-specific variants (NSVs); red, NSVs belonging to a SuSiE credible set (NSV in CS). Variants overlapping open chromatin regions identified by ATAC-seq are highlighted with a black outline. The vertical dashed line marks a |Δsvm| threshold of 2.0179, above which variants are predicted to have a putative effect on chromatin accessibility. Representative NSVs exceeding this threshold or overlapping ATAC-seq peaks are labelled with their rsID.*

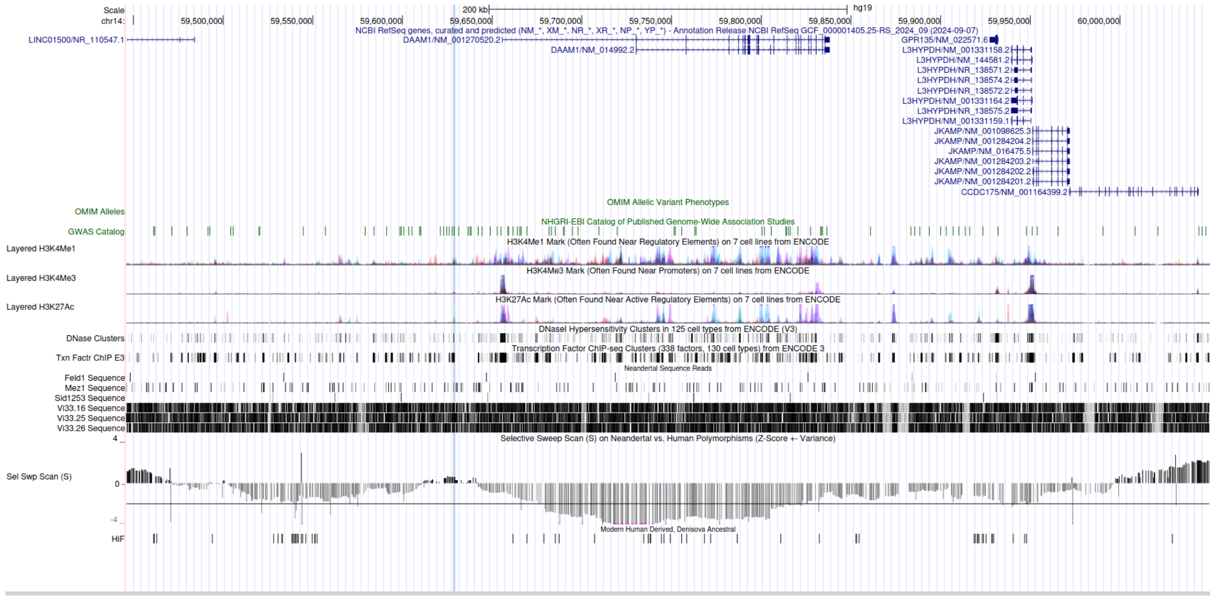


***Figure S4. DAAM1 locus on UCSC genome browser.*** *Several tracks are shown, the blue bar highlights the position of the credible set. Of note, the Sel Swp Scan track shows the S score for positive selection in humans as described in Green et al.*[1] *The large negative region downstream of the credible set is among the top 5% most negative across the human genome (Green et al, table S37). A negative score indicates fewer derived alleles in Neandertal than expected, and may indicate an episode of positive selection in early humans. This score was computed on 5 AMH from the Human Genome Diversity project, including HGDP00521. This sample carries the introgressed allele at rs74826997, suggesting that the local disruption in the selection signal is caused by the introgressed haplotype, and that without it this signal of Neanderthal-AMH divergence could be one of the strongest in the human genome.*

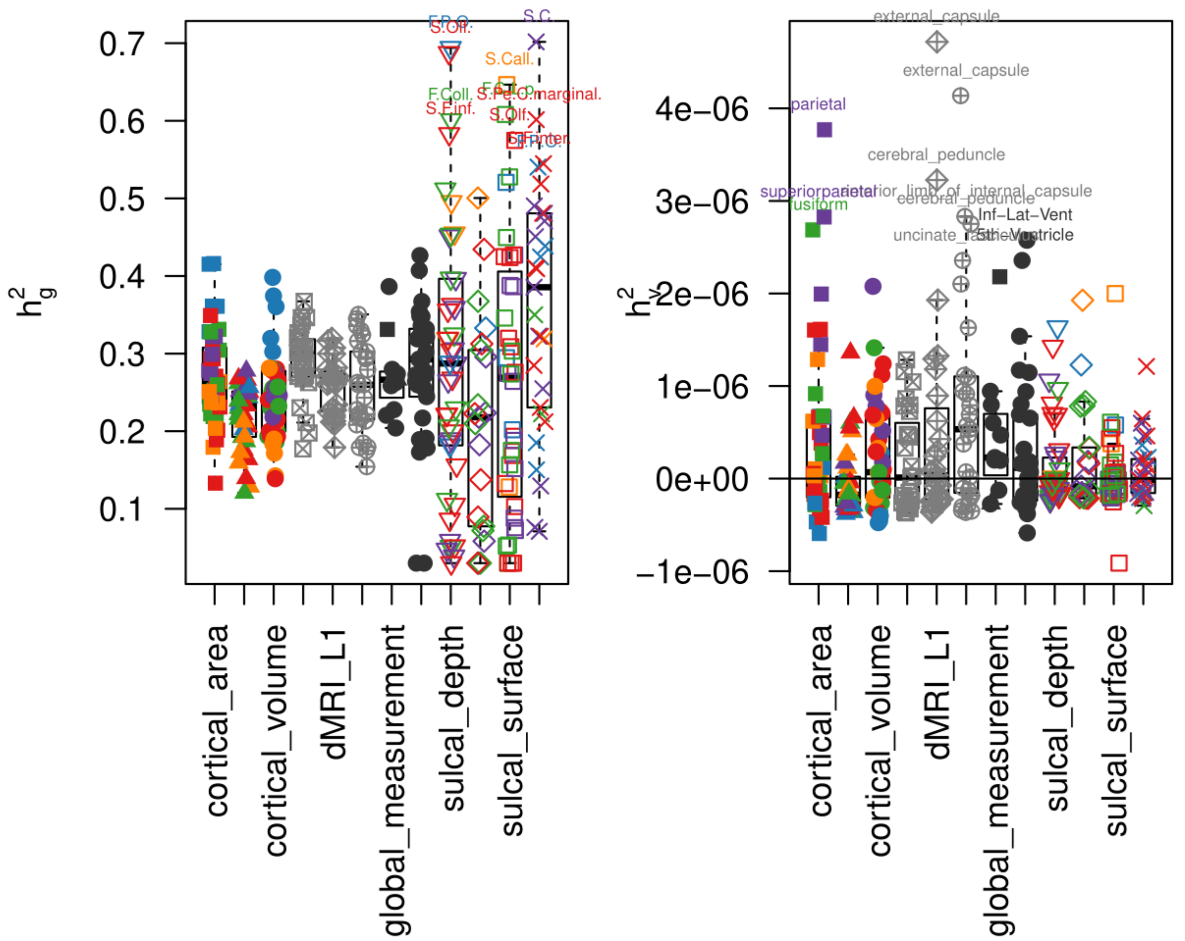


***Figure S5. SLDP heritability estimates. a)*** *total SNP-based heritability for all traits, separated by trait category.* ***b)*** *SNP-based heritability explained by $\Delta p_{Nea,YRI}$ annotation.*

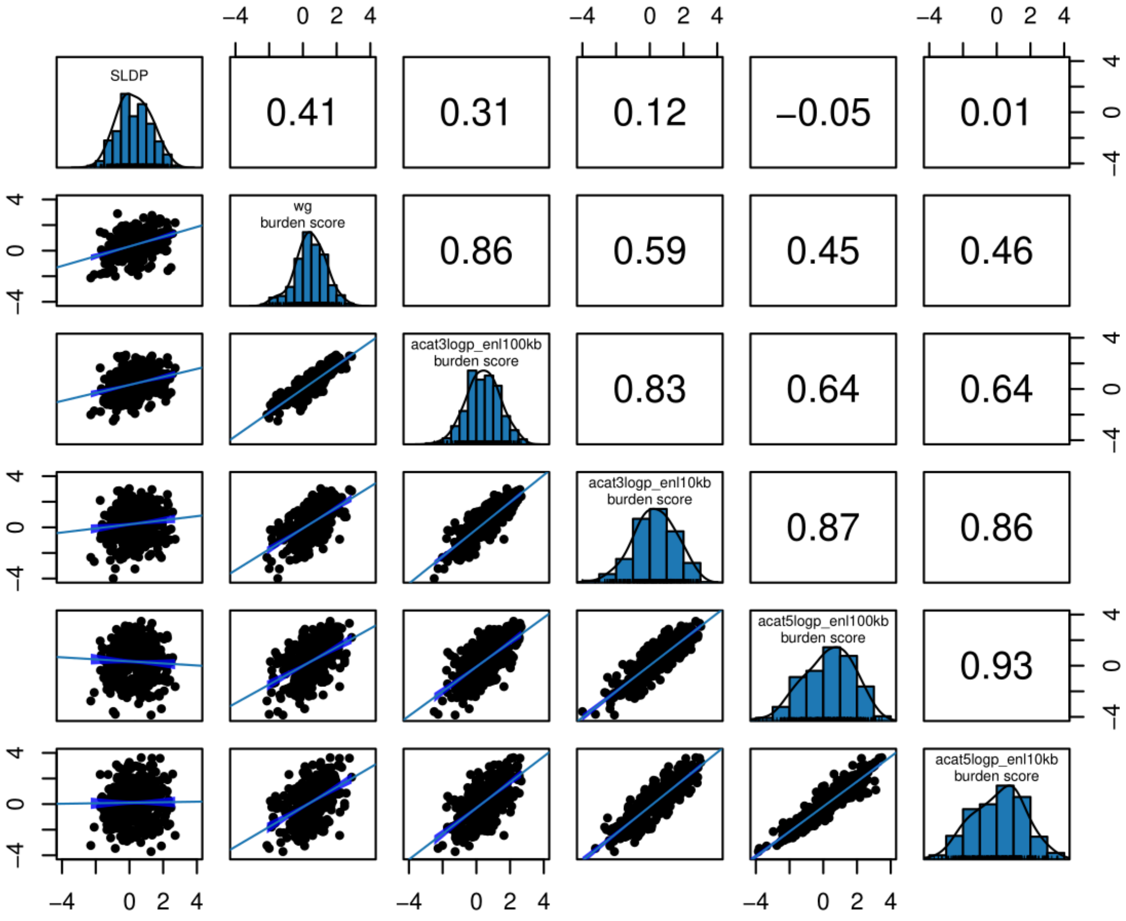


***Figure S6. Correlations across genomic directional effects of Neandertal-derived variants.*** *For different methods (SLDP and regression on Neanderthal burden score) and for different genomic regions on which the score was computed (* $P_{ACAT} < 10^{-5}$ *and* $P_{ACAT} < 10^{-3}$ *, defining 200 or 20 kb loci around significant variants).*

### Supplementary Tables

***Supplementary Table 1. Phenotypes Table.*** *Neuroimaging BIDPs defined by their region of interest (ROI), measurement type, cortical lobe and general category. Label corresponds to the phenotype identifier used in analysis scripts; ROI denotes the anatomical region abbreviation; anatomical_nomenclature provides the full anatomical name where available; measurement specifies the imaging-derived metric; cortical_lobe indicates the broad anatomical grouping; and category denotes the phenotype class.*

***Supplementary Table 2. Loci associated with BIDPs with at least one significant NSV.*** *Each row represents a locus identified through aggregate association testing (ACAT) that harbours at least one genome-wide significant NSV. Chrom, Start, and End indicate the genomic coordinates (hg19) of the locus; ACAT P-value reports the combined p-value summarising association evidence across phenotypes within the locus; Associated Traits lists the brain imaging-derived phenotypes significantly associated with the locus; NSV_in_credible_set and Total_variants_in_credible_set report the number of NSVs and the total number of variants within the credible set, respectively; and NSV_fraction denotes the proportion of credible set variants that are NSVs.*

***Supplementary Table 3. BIDP Colocalization.*** *Full results for the colocalization analysis in each of the 28 loci with significant NSV. The table reports the top trait against which colocalization was performed and the number of colocalization tests. As the colocalization was run with the susie-coloc framework, in some cases susie did not identify a credible set, so the colocalization could not be run. n_H4>H3: number of colocalizations defined as situations where H4 had the highest posterior probability and specifically higher than H3; frac_H4>H3: fraction over total tested ; frac_sum(H3H4)>0.5 ; fraction of possible colocalizations, where the sum of H3 and H4 was above 50% posterior probability; n_CS_avg: average count of independent credible sets identified; NSV>50%_CS: whether NSV were the majority of variants in credible set.*

***Supplementary Table 4. Full Data NSV significant loci.*** *Each row corresponds to a variant (rsID) belonging to one of the 28 associated genomic loci. Genomic coordinates in hg19 and rsID are reported; pval reports the combined p-value (ACAT P-value); awas_pval reports the Neanderthal Association Dosage p-value; ancestry indicates variant as non-Neanderthal (0), introgressed (1) or Neanderthal specific variant (2); and credible_set denotes credible set membership. Columns prefixed ATAC.seq_ contain TRUE/FALSE flags indicating whether the variant overlaps a brain open chromatin region in the corresponding cell type. Cell-type-specific gkm-SVM delta scores are reported for astrocytes (Astro), microglia (Micro), upper-layer excitatory neurons (L2_3), layer 4 excitatory neurons (L4), deep-layer excitatory neurons (L5_6), CGE-derived interneurons (CGE), MGE-derived interneurons (MGE), oligodendrocytes (Oligo), and vascular cells (Vas). Region specifies the genomic window used to identify nearby genes to the variant, and genes_nearby_30000 lists genes located within 30 kb of the variant.*

***Supplementary Table 5. Mental Health GWAS.*** *Category indicates the broad phenotypic category; source identifies the data source or consortium; reference reports the primary publication DOI.*

***Supplementary Table 6. Non-BIDP phenotypes.*** *Traits regressed on the Neanderthal burden score, reporting UKBB field ID and adopted encoding in the note column.*

***Supplementary Table 7. SLDP results.*** *Complete SLDP results for BIDPs and 17 neuropsychiatric and behavioural traits. q-values have been computed separately for these two categories with a Benjamini-Hochberg false discovery rate procedure.*

***Supplementary Table 8. Neanderthal Burden results.*** *Complete results of the neanderthal burden regression for BIDPs and non-BIDP traits. q-values have been computed separately for these two categories with a Benjamini-Hochberg false discovery rate procedure. The first four columns list the results obtained when computing the score on the whole genome (wg), while the second four are the results when computing the score on suggestive loci (suggloci).*

***Supplementary Table 9. Covariates Table.*** *Covariates used in BIDPs GWAS, Neanderthal dosage association and Neanderthal burden regression. The columns #Confs and #Processed indicate the number of unique confound variables and the resulting number from post site-interaction processing, respectively. Processing indicates whether each covariate was treated as continuous (Quantitative) or categorical (Qualitative).*

### Supplementary Notes

### Supplementary Note 1. Partitioned heritability

We tested the application of RHE-mc software from Pazokitoroudi et al.[2] on a sample of 27 traits already examined by Wei et al.[3] to estimate the difference in power attributable to a reduced sample size comparable to the sample size available for BIDPs. RHE-mc is a randomized multi-component Haseman–Elston regression, able to partition phenotype heritability measured in large sample sizes. Following the implementation in Wei et al.[3], to account for the difference in MAF and LD score distributions between NSV and modern human (MH) variants, we assigned the variants to 50 categories by their a) MAF quintile b) LD score quintile and c) NSV vs MH status. We then applied RHE-mc with Nea/MH + MAF + LD annotations to analyze 27 traits, including the top 20 genetic Principal Components estimated from common SNPs, sex, and age as covariates. This analysis was performed first on a set of 337,383 unrelated individuals of European ancestry, then repeated in a random subset of 38,406 individuals, matching BIDP sample size. The heritability estimates were summed across categories defined by MAF and LD score, separately for Nea and MH categories:

$$\widehat{h^2}_{j\in\{Nea,MH\}} = \sum_i \widehat{h^2}_{i,j}$$

Standard error was computed using standard error propagation assuming independent h2 estimates across SNP categories. When comparing our estimates with those reported by Wei et al.[2] we did not notice a directional bias (**Figure S7A**). Standard errors were similar for the larger sample set (median ratio = 0.964) but showed marked increase in the analysis with the BIDP-comparable sample set: median 3.18-fold increase for MH and 4.28-fold increase for NIM h2 estimates (**Figure 7B,C**). This is comparable or higher than the theoretically expected increase in estimation standard error due to the decrease in sample size (2.96-fold for 11.4% of sample size). As just 6 traits showed significant heritability mediated by NSV in Wei et al., and as that would result in about a three-fold decrease in the Z-statistic, we deemed unlikely for this approach to find any significant heritability contribution from Neanderthal in our data.

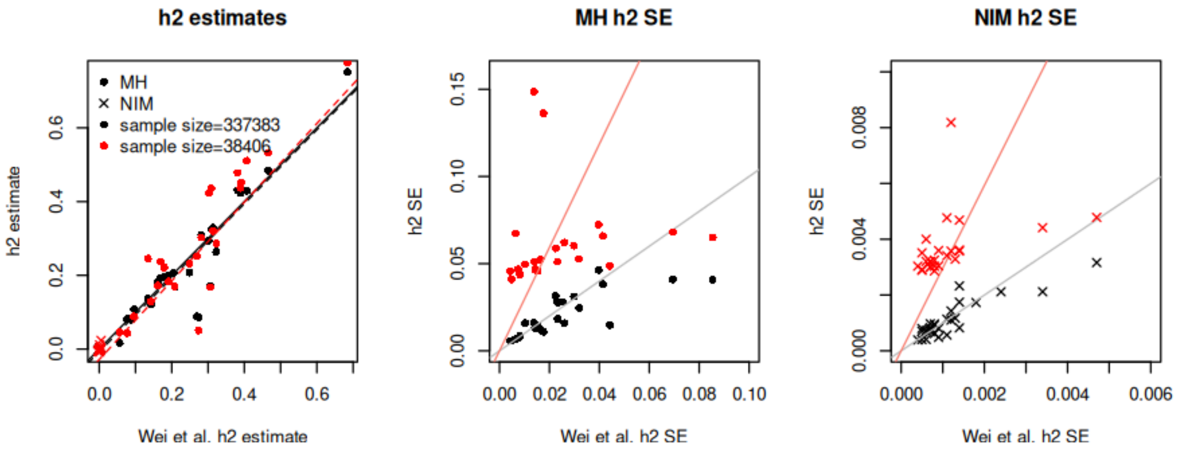


***Figure S7 - RHE-mc results on a subset of the UK Biobank.*** *a) heritability estimates compared with the estimates reported by Wei et al. Red dots indicate estimates obtained with a sample size comparable to our BIDPs analysis, crosses indicate estimates for the heritability explained by Neanderthal-specific variants, termed "NIM" by Wei et al. b) and c) standard errors of the estimate separately for modern human variants and for NIM, respectively.*

### Supplementary Note 2 - Significant loci mediated by NSV

##### chr1:59217533-59432116 - JUN

NSVs in the credible set cluster around 59.25-59.35 Mb, and overlap open chromatin across all brain ATAC-seq cell types (Astro, CGE, L2_3, L4, L5_6, MGE, Micro, Oligo and Vas) with a subset exceeding the deltaSVM threshold for putative chromatin accessibility alteration in Microglia. None of these functionally annotated variants is a significant eQTL or sQTL in GTEx, but the closest gene, JUN, has a prominent neurodevelopmental and neuropsychiatric literature, making it a likely candidate in mediating the effect of this locus.

In humans, JUN expression is dysregulated in the post-mortem prefrontal cortex of individuals with major depressive disorder and bipolar disorder, implicating AP-1 signalling in the pathophysiology of affective illness[4]. Genome-wide transcriptomic studies of human brain tissue have further identified AP-1 complex members -including JUN- as convergent hubs of gene co-expression networks disrupted across schizophrenia, bipolar disorder, and autism spectrum disorder[5]. In the context of neurodevelopment, JUN has been identified as a downstream target of disrupted-in-schizophrenia 1 (DISC1) signalling, linking AP-1-mediated transcription to cortical progenitor proliferation and the aetiology of neurodevelopmental psychiatric conditions in humans[6]. This locus is associated with 2 traits, specifically posterior corona radiata mean diffusivity (MD) and posterior thalamic radiation fractional anisotropy (FA). Both measures reflect water diffusion measurements of white matter tracts connecting posterior cortical regions to more caudal structures such as the brainstem and the thalamus, respectively.

##### chr3:190491314-190778438 - GMNC

Multiple variants falling between 190.55-190.70 Mb are NSVs belonging to a credible set and overlap brain ATAC-seq regions across most cell types (Astro, CGE, L2_3, L4, L5_6, MGE and Oligo) with a subset exceeding the deltaSVM threshold for putative chromatin accessibility alteration in Microglia, Oligodendrocytes and MGE. None of these functionally annotated variants is a significant eQTL or sQTL in GTEx; however, the proximity to GMNC (Geminin Coiled-Coil Domain Containing, also known as GEMC1), a transcriptional regulator of multiciliated cell (MCC) fate[7], implicates GMNC as a plausible candidate gene in mediating the effect of this locus. GMNC and its paralog MCIDAS are key regulators of multiciliated ependymal cell generation in the brain, operating upstream of the FOXJ1 and c-Myb transcription factors, a relationship conserved across vertebrates[8]. In humans, loss-of-function variants in *MCIDAS* and *FOXJ1* cause primary ciliary dyskinesia and are directly associated with congenital hydrocephalus, highlighting the critical role of this transcriptional programme in human ventricular development and cerebrospinal fluid (CSF) homeostasis[9]. This locus is significantly associated with 36 traits, spanning ventricular volumes (3rd, 4th, and lateral ventricles, inferior lateral ventricle, and choroid-ventricular volume), choroid plexus volume, brainstem volume, and global brain segmentation metrics. Associations also extend to sulcal morphology measures reflecting periventricular sulcal opening patterns, cortical regions anatomically in proximity to the ventricular system - including entorhinal, parahippocampal, precuneus, superior parietal, and lateral occipital area and volume - and subcortical structures including the amygdala and nucleus accumbens. White matter associations encompass the fornix (FA, L1, and MD), fornix cres and stria terminalis FA, tapetum L1, anterior corona radiata FA, posterior limb of the internal capsule L1, and corpus callosum segments (central and mid-anterior).

##### chr4:121446341-121865788 - PRDM5

NSVs within the credible set span 121.5-121.8 Mb and overlap brain ATAC-seq regions across most cell types (Astro, CGE, L2_3, L4, L5_6, MGE, Oligo and Vas) with a subset exceeding the deltaSVM threshold for putative chromatin accessibility alteration in L5_6 and L4 neurons. Among the several variants overlapping brain ATAC-seq peaks, rs11733936, rs13121457, rs17412912, rs2290, rs34827843, rs4473670, rs4555666, and rs72676317 act as eQTLs (GTEx), while rs11721741, rs12508048, rs17051266, rs4833686, rs71600496, and rs76291603 are annotated as both eQTLs and sQTLs chiefly for the gene *PRDM5*. Two variants exceed the top 1% deltaSVM threshold derived from gkm-SVM models in L5_6 and L4 neurons: rs35013646, an eQTL linked to *PRDM5*, and rs3804169, which functions as both an eQTL and sQTL for the same gene. *PRDM5* is a transcription factor which has been shown to regulate Wnt/β-catenin signaling in zebrafish[10], a pathway critically involved in neural progenitor proliferation, cortical development, and ventricular zone organisation across vertebrates. Finally
This locus is significantly associated with 23 traits, with a dominant signature of altered cortical thickness spanning frontal and parietal regions. Thickness associations encompass global mean cortical thickness and a broad frontoparietal network including superior frontal, rostral and caudal middle frontal, lateral and medial orbitofrontal, pars opercularis, pars triangularis, precentral, paracentral, inferior parietal, superior parietal, supramarginal, and precuneus cortex, as well as entorhinal area and middle temporal volume. White matter associations include the superior longitudinal fasciculus fractional anisotropy (FA) and anterior corona radiata FA, alongside sulcal morphology of the lateral anterior occipito-temporal sulcus (surface area, mean depth and hull junction length).

##### chr5:55908915-56283743 - SETD9/MAP3K1

NSVs belonging to the credible set are distributed across 55.95-56.20 Mb and overlap with all brain ATAC-seq cell types (Astro, CGE, L2_3, L4, L5_6, MGE, Micro, Oligo and Vas)(refer to Figure S3). Among variants overlapping brain ATAC-seq peaks, rs12659430, rs1862620, rs62355881, rs62355907, rs62355943, and rs72758038 are annotated as both eQTLs and sQTLs (GTEx) for two previously uncharacterized genes (ENSG00000225940 and ENSG00000271828) and SETD9, albeit with lower significance. None of these variants exceed the top 1% deltaSVM threshold derived from gkm-SVM sequence-based modelling of chromatin accessibility. While for these genes the implication in neurodevelopment is not evident, the credible set also overlaps MAP3K1, which encodes a serine/threonine kinase upstream of the JNK and ERK-MAPK pathways, signalling pathways that play central roles in neurodevelopment[11,12].This locus is significantly associated with 18 traits, including global mean cortical thickness, brainstem volume, and a constellation of regional cortical thickness and volume measures. Thickness associations span the precuneus, superior parietal, posterior cingulate, parahippocampal, fusiform, lingual, rostral and caudal middle frontal cortices, alongside composite regional metrics for cingulate, frontal, parietal, and temporal thickness. Volume associations include the precuneus, superior parietal, lingual, and cingulate and parietal lobes.

**chr6:99727313-100239801 - TSTD3**

NSVs in the credible set span approximately 99.8-100.1 Mb and overlap with all brain ATAC-seq cell types (Astro, CGE, L2_3, L4, L5_6, MGE, Micro, Oligo and Vas) with a subset exceeding the deltaSVM threshold for putative chromatin accessibility alteration in L5_6 cells (Figure S3). Among variants overlapping brain ATAC-seq peaks, rs330872, rs491178, and rs609669 are annotated as both eQTLs and sQTLs (GTEx). Furthermore, one variant (rs12529031) exceeds the top 1% deltaSVM threshold derived from gkm-SVM sequence-based modelling of chromatin accessibility, annotated as eQTL and sQTL for a large set of genes including, in order of decreasing significance: TSTD3, USP45, PNISR-AS1, CCNC, FAXC, COQ3, PNISR. Of these, the expression of the first three genes is associated with this variant in brain tissues, but they do not seem to have any obvious connections with neurodevelopmental phenotypes. This locus is significantly associated with 1 trait: cerebellar cortex volume.

**chr6:133465734-133681377 - EYA4**

NSVs in the credible set span around 133.5-133.6 Mb and overlap brain ATAC-seq in several cell types (CGE, L2_3, L4, L5_6, MGE and Oligo). Among these, one variant, rs9375942, exceeds the deltaSVM threshold and is predicted to alter chromatin accessibility in vascular cells. This locus is significantly associated with 12 traits spanning cortical morphology, total brain volume and white matter microstructure. rs9375942 lies in a cis-regulatory region with significant eQTL and sQTL effects on EYA4 in multiple human brain tissues, including frontal cortex and anterior cingulate (GTEx). A member of the Eyes Absent transcriptional coactivators, EYA4, when mutated, is responsible for postlingual, progressive, autosomal dominant hearing loss[13], so the connection of this gene with the several brain morphology associations found here is not clear. Total surface, caudal middle frontal area, parstriangularis volume, and cingulate/frontal area and volume, diffusion measures of the corticospinal tract and pontine crossing tract (L1, MD) were found as associated in our data.

**chr20:45086396-45304763 - SLC13A3**

NSVs in the credible set cluster across 45.10-45.28 Mb and overlap with all brain ATAC-seq cell types (Astro, CGE, L2_3, L4, L5_6, MGE, Micro, Oligo and Vas). Fine-mapping of the credible set reveals functional regulatory evidence for several NSVs (refer to Figure S3). Among variants overlapping brain ATAC-seq peaks, rs2694876, rs3091661, rs3092554, rs386843, rs393990, rs399940, rs430891, rs432915, rs439143, rs454103, and rs847068 are all annotated as both eQTLs and sQTLs linked to *OCSTAMP* and *SLC13A3* (GTEx). rs439143 and rs863674 at this locus have been found in significant excess of neanderthal allele homozygosity in Morez Jacobs et al (Table S6)[14], therefore suggesting that this locus might be under adaptive selection of the introgressed haplotype. In the human brain, SLC13A3 is expressed predominantly in astrocytes. Variants in SLC13A3 have been associated with leukoencephalopathy with episodic neurological crises, highlighting its importance in white matter integrity[15]. This locus is significantly associated with 3 traits, all reflecting white matter microstructure of frontoparietal association and projection fibres. Associations include the superior corona radiata MD and the superior longitudinal fasciculus FA and MD.

**chr22:40445796-40820963 - TNRC6B**

The credible set at this locus is composed for the 41% by NSVs: although not strictly qualifying as a NSV-mediated locus, we discuss it briefly as a suggestive case. NSVs in the credible set cluster around 40.55-40.70 Mb and overlap with all brain ATAC-seq cell types (Astro, CGE, L2_3, L4, L5_6, MGE, Micro, Oligo and Vas).

Among variants overlapping brain ATAC-seq peaks, rs138039, rs5750911, rs5995839, and rs8141661 are all annotated as both eQTLs and sQTLs linked to TNRC6B (GTEx). rs5750911 has been previously associated with smoking initiation in large-scale GWAS, raising the possibility that this variant may influence addiction-relevant neurobiology through transcriptional and splicing regulation of TNRC6B in addition to its effect on brain morphology. Rare human variants in TNRC6B have been associated with neurodevelopmental delay and autism spectrum disorder in recent sequencing studies, consistent with its central role in post-transcriptional gene regulation in the developing human brain[16]. This locus is significantly associated with two traits: insular cortex volume and total cerebral white matter volume.
